# Interaction of the Tau fibrils with the neuronal membrane^†^

**DOI:** 10.1101/2022.12.14.520516

**Authors:** Unmesh D. Chowdhury, Arnav Paul, B.L. Bhargava

**Author notes:** Electronic Supplementary Information (ESI) available: [The Supplementary Information comprises of additional simulation details in Section 1, RMSD of the tau fibrils for the 1 *μ*s simulations (Figure S1), initial and final configuration of the neuronal membrane (Figure S2), RMSD (Figures S4 – S5), RMSF (Figure S6 – S7), tables containing the radius of gyration values (Table S2 – S3), tables of the SASA values (Table S4 – S5) and secondary structure content of the independent simulations (Table S6 – S7), approach distance of the tau fibril over the neuronal membrane (Figure S8), snapshot of positively charged residues over the bilayer plane (Figure S9), snapshots of final configurations of PHF and SF structures (Figure S10), bilayer properties (Table S8) and two dimensional bilayer thickness for the independent simulations (Figure S11 – S12), cholesterol tilt angle distribution of the systems (Figure S13).].

## Abstract

Tau proteins are gaining a lot of interest recently due to their active role in causing a range of tauopathies. Molecular mechanisms underlying the tau interaction with the neuronal membrane are hitherto unknown and difficult to characterize using conventional experimental methods. Starting from the cryo-EM structure of the tau fibrils, we have used atomistic molecular dynamics simulations to model the interaction between the fibril and neuronal membrane, with explicit solvation. The dynamics and structural characteristics of the tau fibril with the neuronal membrane are compared to the tau fibril in the aqueous phase to corroborate the effect of the neuronal membrane on the tau structure. The tau fibrils are in general more compact in the presence of neuronal membrane compared to their structure in the water medium. We find that the number of *β* -sheet residues of the tau fibrils are different in the case of two polymorphs, paired helical filament and straight filaments (PHF and SF) in the two media. PHF is found to approach closer to the neuronal membrane than the SF. The negatively charged lipids in the neuronal membrane are found to mediate the tau-neuronal membrane binding. Our study initiates the understanding of tau conformational ensemble in the presence of neuronal membrane and sheds light on the significant tau – membrane interactions. The simulation times of our report might limit the conformational sampling required to observe membrane permeation, nevertheless it provides significant insights into fibril – neuronal membrane interactions.

## 1 Introduction

The tau protein plays an important role in microtubule stabilization and aggregation^1^. In diseased neurons, tau loses the capacity for microtubule binding and forms toxic aggregates^2^. These toxic aggregates are the causative agents for a series of neurodegenerative diseases collectively known as tauopathies^3^. Tauopathy comprises a set of neurodegenerative diseases like Alzheimer’s Disease, Pick’s disease, frontotemporal lobar degeneration, etc. Hyperphosphorylated tau has a positive effect on the progression of Alzheimer’s disease (AD). Tau is an intrinsically disordered protein (IDP) that shows a diverse conformational ensemble^4^. Tau and the other disease-forming aggregates have been shown to evolve through the liquid – solid and liquid – liquid phase transition of the proteins^5^. The conformational dynamics of the tau fibrils has a significant role in the pathogenesis of tauopathy as shown from the earlier molecular dynamics studies^6–13^. Owing to the failure of the amyloid-beta as the drug-binding target for Alzheimer’s disease (AD), the current focus is mostly on the tau fibrils and their underlying conformational changes in regulating the concurrent tauopathies^14,15^. The interaction of the lipid bilayers with the tau is pivotal in understanding the mechanism of toxicity of the tau fibrils^16^. Our earlier MD study was focused on the effect of the net charge of the lipids and the effect of cholesterol in determining the tau conformational changes^17^. It has been widely accepted that the negatively charged lipids influence the tau aggregation^18,19^.

The emergence of single molecular techniques has further enhanced our knowledge related to tau aggregation and conformational states of IDPs^20^. Single-molecule Fluorescence resonance energy transfer (smFRET) and fluorescence correlation spectroscopy (FCS) have shown that the tau proteins bind to the DMPS membrane in a concentration dependent manner. At low DMPS concentrations the tau proteins form oligomers. At high DMPS concentration the tau proteins inhibit amyloid formation due to the stable binding with DMPS^21^. Atomic Force Microscopy (AFM) study has explained the effect of PIP_2_ lipids on the Tau microtubule-binding construct K18^22^. Intrinsically disordered proteins (IDPs) have also been characterized using molecular dynamics simulations with the emergence of the optimized force fields and compatible water models^4,23–25^. The dynamical changes in the IDPs are experimentally derived using the small angle neutron/X-ray scattering (SANS/SAXS), nuclear magnetic resonance (NMR), circular dichroism (CD), fluorescence resonance energy transfer (FRET), etc which are usually insufficient to obtain complete dynamical information of the IDPs^26–29^. Thus, molecular dynamics simulations have been widely used to characterize the IDPs responsible for AD pathogenesis, especially the amyloid-beta peptide and amyloid fibrils^30–34^. Strodel et al. have studied the interactions of the amyloid-beta dimer with the axonal membrane^35^. In the pursuit of gaining atomic level insights into the interaction between the tau fibril and neuronal membrane, we have modeled the paired helical filaments (PHF) and straight filaments (SF) in the presence of model neuronal membrane for a cumulative sampling of 4 microseconds. To the best of our knowledge, this is the first comprehensive computational work on the tau fibrils in presence of the neuronal membrane.

The starting configuration of the lipid bilayer along with the tau fibril polymorphs are shown in Figure 1. The proportion of the individual lipid molecules is shown as a pie chart in Figure 1(b). We have maintained the same starting configurations of the tau fibril over the neuronal membrane. We elucidate the differences in the tau morphology in the solution phase and the neuronal membrane from the atomistic simulations. The computational modeling presented here only accounts for the early stages of the tau interaction with the neuronal membrane. The entire event of the tau permeation through the neuronal membrane is beyond the scope of this study due to the limitations of the computational resources required for all-atom simulations using explicit solvation.

**Fig. 1.**
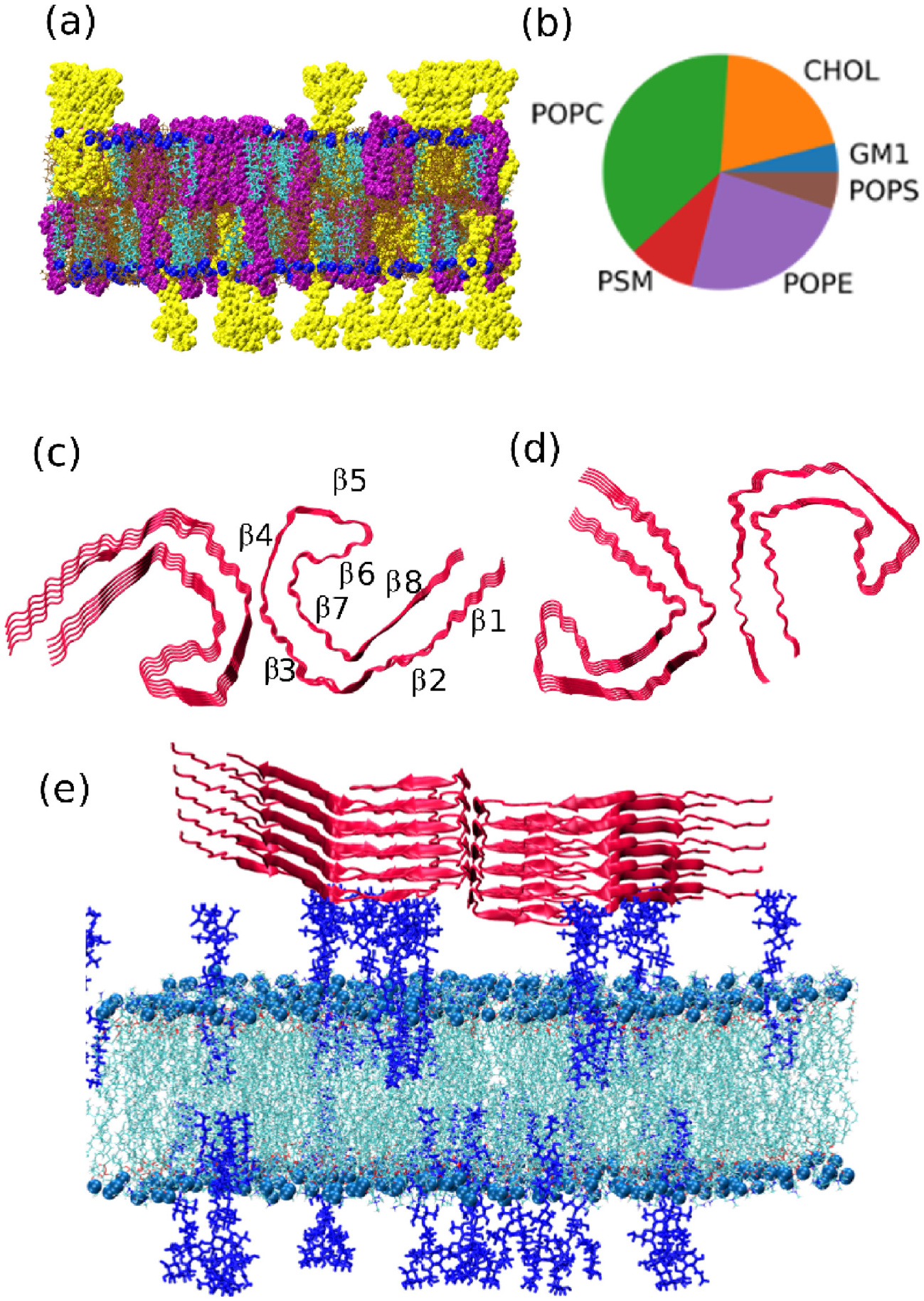
(a) The neuronal membrane with the phosphorus atoms, GM1 and PSM shown as blue, yellow and purple vdw spheres respectively, and cholesterol and other lipids shown as cyan and brown lines respectively. (b) the pie chart showing the composition of different lipids in the neuronal membrane, (c) the PHF structure shown in new cartoon representation with the different zones of *β* -sheet regions marked, (d) the SF structure, (e) the starting configuration of the PHF fibril above the neuronal membrane, where the GM1 lipids are shown as blue licorice, phosphorus atoms and other lipids are shown in blue and cyan respectively.

## 2 Methodology and simulation details

The systems modeled are composed of the PHF and SF tau fibrils taken from the PDB id 503L and 503T respectively, and were simulated in the aqueous phase and the neuronal membrane^36^. The cryo-EM structures are having a resolution of 3.4 – 3.5 Å and are derived from the brain of an individual having Alzheimer’s disease. The SF and PHF fibrils are having eight zones of *β* -sheet regions with similar protofilament cores. The amino acid residues in the R3-R4 stretch are 306VQIVYKPVDLSKVTSKCGSLGNIHHKPGGGQ336VEVKSEKLDFKD RVQSKIGSLDNITHVPGGGN368KKIETHKLTF378. The difference in the structure of PHF and SF stems from the difference in the lateral contacts at the dimer interface.

Hence, computational modeling of the dimer tau fibrils become important to account for the tau polymorphism. The PHF and SF fibrils are also simulated in the water box to compare with the neuronal membrane simulations. The neuronal membrane comprises 38% 1-palmitoyl-2-oleoyl-sn-glycero-3-phosphocholine (POPC), 24% 1-palmitoyl-2-oleoyl-sn-glycero-3-phosphoethanolamine (POPE), 5% 1-palmitoyl-2-oleoyl-snglycero-3-phospho-L-serine (POPS), 20% cholesterol (CHOL), 9% palmitoylsphingomyelin (PSM) and 4% monosialotetrahexosylganglioside (GM1). The tau fibrils are modeled using the CHARMM-36m force field which is shown to perform best for the IDPs^37^. The neuronal membrane composition of symmetric lipid bilayers in the PHF structure is 247 POPC, 156 POPE, 130 cholesterol, 60 PSM, 33 POPS, and 26 GM1 molecules. The corresponding symmetric bilayers in the case of SF structure comprise of 209 POPC, 132 POPE, 110 cholesterol, 49 PSM, 28 POPS, and 22 GM1 molecules. The bilayers are generated using the CHARMM-GUI membrane builder^38^. The membranes are comprised of layers of water molecules above and beneath the upper leaflet and lower leaflet of the bilayer.

The fibrils are modeled using the CHARMM-36m force field and the lipids are modeled using the CHARMM-36 force field^39,40^. The all-atom simulations have been performed using the GRO-MACS molecular dynamics software (version -5.1.4)^41^. The systems were initially energy minimized using the steepest descent algorithm to negate the unfavorable contacts. This was followed by the six steps of equilibration as prescribed in the default CHARMM-GUI input generation. Finally, systems were equilibrated in the NPT ensemble and the production runs were carried out for 1 *μ*s. The pressure is maintained using the semi-isotropic Parrinello-Rahman pressure coupling scheme^42^. The temperature is maintained using the Nosé-Hoover thermostat^43^. The periodic boundary condition (PBC) is applied in all directions. Particle Mesh Ewald (PME) is used to model long-range electrostatics^44^. The van der Waals and coulombic cutoff are set to 1.2 nm. The van der Waals interactions are modeled using the force switch algorithm. All the bonds involving hydrogen atoms were restrained using the LINCS algorithm^45^. Water and chloride ions were used to dissolve and neutralize the protein-lipid systems. The water molecules have been modeled using the TIP3P water model^46^. Additional 150 mM of KCl was added to model the physiological salt concentration. Two additional simulations of 100 ns each have been performed in case of both the PHF and SF systems. Control simulations of 500 ns were performed in the water phase with PHF and SF. These simulations are also complemented with two independent simulations of 100 ns each.

Similarly, neuronal membrane without the tau fibril is simulated taking 200 lipids in total. The pure neuronal membrane simulations have been performed for 500 ns each with two independent starting configurations. Further details of the simulated systems are given in Table 1.

**Table 1.**
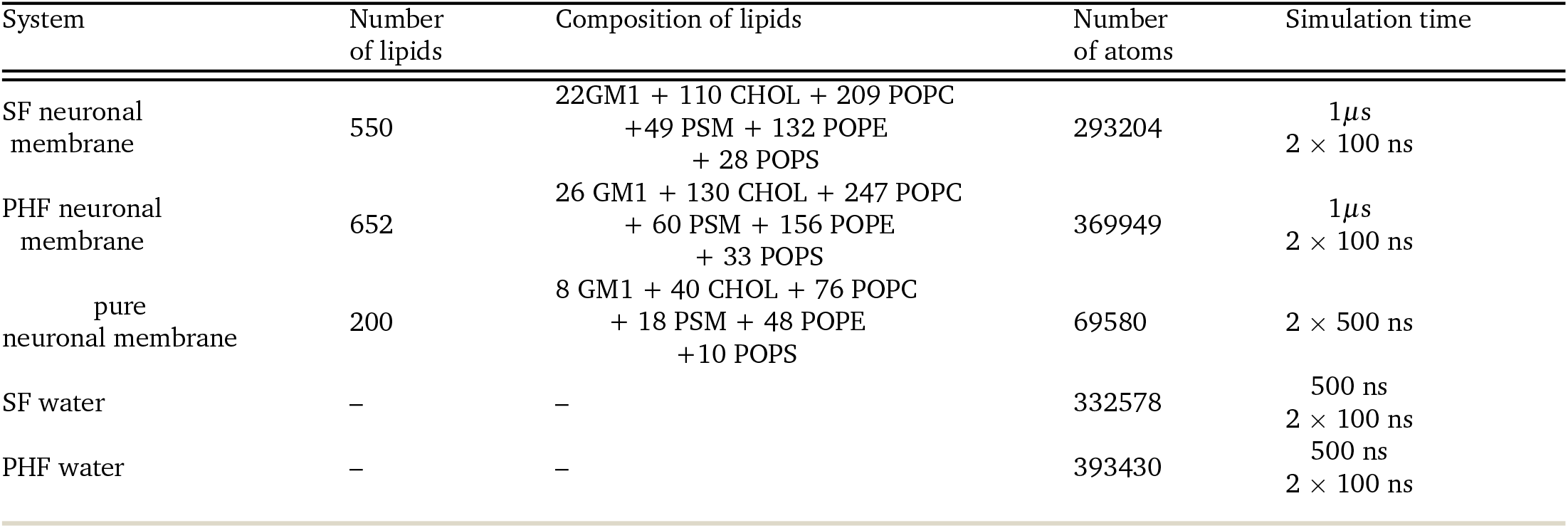
The details of the simulated systems

| System | Number of lipids | Composition of lipids | Number of atoms | Simulation time |
| --- | --- | --- | --- | --- |
| SF neuronal membrane | 550 | 22GM1 + 110 CHOL + 209 POPC<br>+ 49 PSM + 132 POPE<br>+ 28 POPS | 293204 | 1μs<br>2 × 100 ns |
| PHF neuronal membrane | 652 | 26 GM1 + 130 CHOL + 247 POPC<br>+ 60 PSM + 156 POPE<br>+ 33 POPS | 369949 | 1μs<br>2 × 100 ns |
| pure neuronal membrane | 200 | 8 GM1 + 40 CHOL + 76 POPC<br>+ 18 PSM + 48 POPE<br>+ 10 POPS | 69580 | 2 × 500 ns |
| SF water | – | – | 332578 | 500 ns<br>2 × 100 ns |
| PHF water | – | – | 393430 | 500 ns<br>2 × 100 ns |

All the analyses have been carried out with the data from the entire span of 1 *μ*s or 500 ns unless mentioned otherwise. Additional simulation details for all the systems (fibril+membrane, fibril+water, membrane only) are included in the Supplementary Information (Section 1). The initial configurations of the tau fibril above the membrane along with the simulation cell are given in Supplementary Information (Figure S1). The snapshot of the neuronal membrane without the tau fibrils at the beginning and at the end of the simulation are also included in the Supplementary Information (Figure S2).

The analysis codes have been written using MDAnalysis python library and *gmx* tools. Clustering analysis was done using the GROMOS algorithm with the RMSD cutoff of 0.2 nm^47^. Bilayer thickness is calculated from the average position (perpendicular to interface) of the phosphate groups in both the leaflets. The area per lipids (APL) is calculated by taking the product of the *x* and *y* dimensions of the simulation box and dividing it by the total number of lipids. The number of contacts are calculated between the tau fibril residues and each of POPC, POPS, POPE, PSM or GM1 lipid. A contact is defined when the distance between any two non-hydrogen atoms from the residue and lipid in question is within 1.0 nm.

## 3 Results and Discussion

### 3.1 Structural Features

The polymorphic tau fibrils show differential stability due to the change in the environment in going from the neuronal membrane to the water medium. The root mean square deviations (RMSDs) of the tau fibrils for the 1 *μ*s simulations are shown in Supplementary Information (Figure S3). SF-neuron and PHF-neuron refers to the neuronal membrane simulations and SF-water, PHF-water refers to the simulations in the water medium. The RMSDs for the structural polymorphs in the independent simulations are included in the Supplementary Information (Figure S4 – S5). To characterize the structural stability, we have calculated root mean square fluctuations (RMSFs) of the C-*α* residues of the tau fibril, which are shown in Figure 2(a). The RMSF for the independent simulations are given in the Supplementary Information (Figure S6 – S7).

**Fig. 2.**
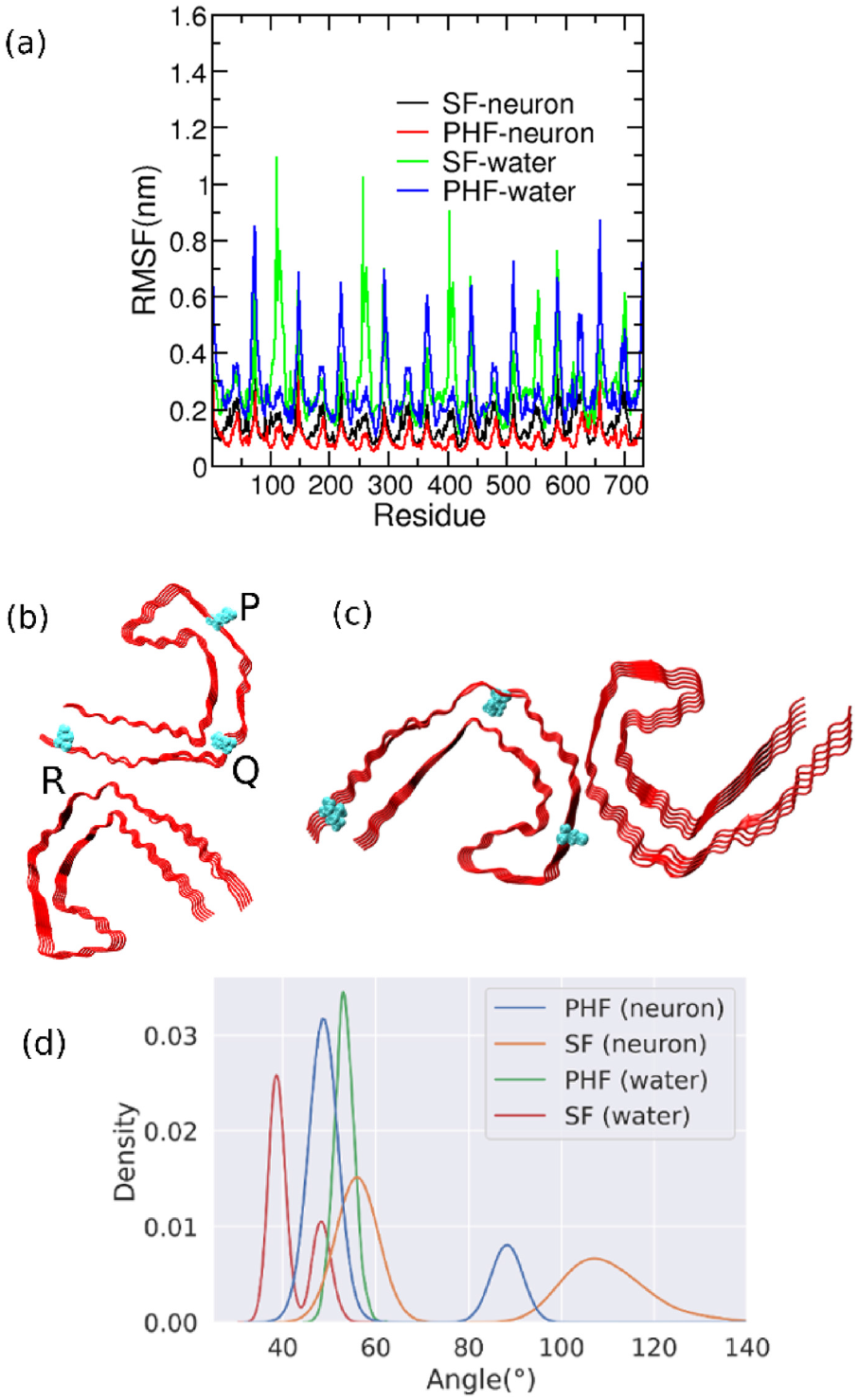
(a) RMSF of the C-*α* atoms of the residues of the tau fibril in the presence of neuronal membrane and in the water medium. (b)-(c) Pictorial representation of the angle of coverage (∠PQR) and (d) the distribution of the angle of coverage in various systems.

The neuronal membrane decreases the structural fluctuations of both the PHF and the SF polymorphs of the tau fibril when compared with the simulations in water. The compaction of the tau proteins in the lipids when compared to the water medium is in accordance with the experimental reports published on the model lipids^48,49^. The RMSF of the PHF is less than that of the SF structure conforming to the fact that the PHF structures are more stable than the SF structures^50^. We have also calculated the angle of coverage, subtended by the three residues as shown in Figure 2(b). The angle defines the characteristic ‘C’ shape of the PHF and SF structures of tau as shown earlier^17^. The most probable angle of coverage for PHF in the water medium is 53°, and in the neuronal membrane it is 57° with a small distribution at 88°. In the neuronal membrane with SF, the angle of coverage is peaked at 56° with a smaller distribution at 106°. In case of SF in water medium, the peak is observed at 39° with a small distribution at 48°. The corresponding angle for the PHF and the SF structures in the cryo-EM structures are 58° and 57° respectively. Thus, the angle of coverage is decreased in the water medium compared to the neuronal membrane and the cryo-EM structure, and this is mostly driven by the higher flexibility of the tau fibrils in the water medium.

The positively charged residues interacting with the lipid bilayers are shown to be important for the fibril-membrane interaction. This has been established in the earlier studies related to the A*β* protofilament^51^, model peptides^52^ and the hIAPP fibrils^31^. The coulombic interactions are found to be important factors for fibril-lipid headgroup binding. It has also been found experimentally that the negatively charged DMPG membranes show differential binding with the charged and acetylated N-terminus of the tau hexapeptide^53^. Also, the attachment of the human tau on the human brain lipid membranes are dependent on the cation present, providing evidence of the electrostatic forces governing the interaction^18^. Sodium (Na+) is found to facilitate the attachment, whereas potassium (K+) inhibits the process. Hence, it has been proven that the coulombic interactions are pivotal to the tau-membrane interactions and the associated toxicity.

The distance profile of the positive charged residues with the P atoms of the bilayer as a function of time are shown in Figure 3. The positively charged residues are shown in blue surface representation. The mean distances of the positively charged PHF residues with the P atoms of the headgroups are shown to be less than that of the SF structures. This is also evident from the approaching depth of the PHF and SF structures over the neuronal membrane, which is shown in Supplementary Information (Figure S8). A different view of the positively charged residues over the bilayer plane is shown in Supplementary Information (Figure S9), where the positively charged residues are viewed from the top of the bilayer.

**Fig. 3.**
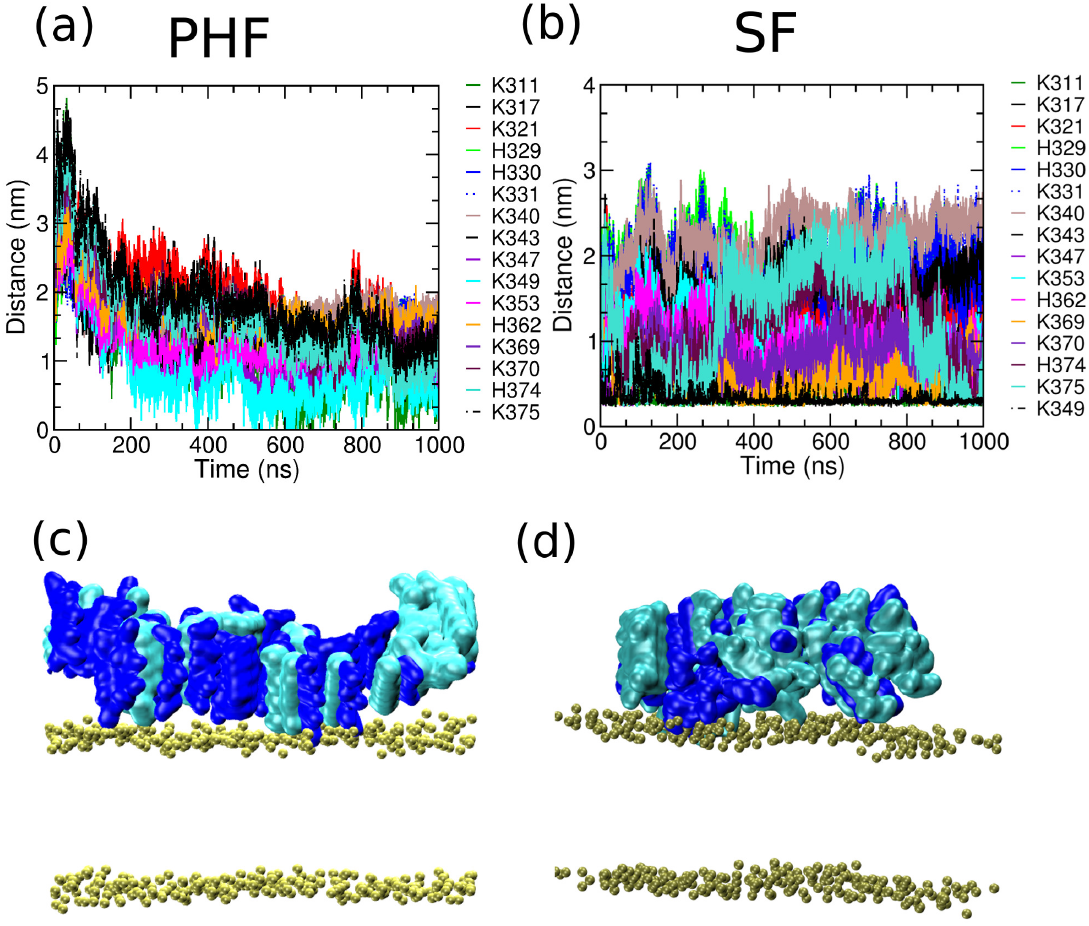
(a)-(b) are the distance profiles of the positively charged residues of the tau fibril with the phosphorus atoms of the lipid head groups. (c)-(d) are the representative tau fibril conformations showing positively charged residues in blue surface representation and other groups in cyan surface representation.

To study fibril membrane interaction, we calculated the twodimensional profiles for the fibril – lipid and intra-fibril contacts with respect to the fibril approach distance, which is shown in Figure 4. The fibril – lipid contacts are normalised with respect to the total number of lipid molecules. The approach distance is calculated as the distance along z-axis between the center of mass of the membrane and that of the fibril. PHF is more closer to the neuronal membrane than the SF structure. The average number of fibril-lipid contacts are found to increase with closer approach of the fibril to the bilayer. The PHF approaches closer to the bilayer than the SF fibril. In the SF structure, the distribution of the fibril-lipid contacts and the approach distance are broader. Thus, we infer that in the timescale of our simulations, PHF has more fibril-lipid contacts, which results in the closer approach to the membrane. PHF also shows a higher number of intra-fibril contacts at a smaller approach distance compared to the SF structure. The higher number of intra-fibril contacts suggests higher stability of PHF structure with respect to the SF structure with the concomitant approach of the tau fibril towards the neuronal membrane.

**Fig. 4.**
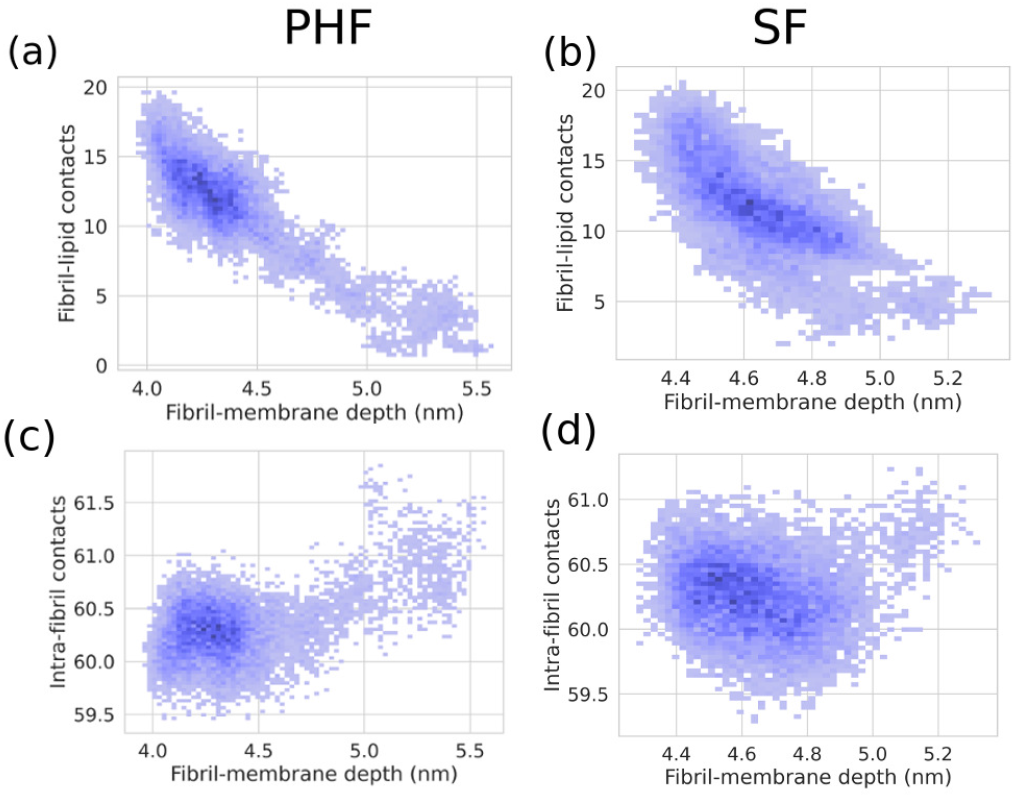
The two dimensional depth profile of the fibril center of mass along the z-axis with respect to the fibril – lipid and intra-fibril contacts. (a) and (c) are the profiles of fibril – lipid and intra-fibril contacts respectively with the approach distance for PHF and (b) and (d) are corresponding graphs for the SF structures.

### 3.2 Radius of Gyration (R*g*) and Solvent Accessible Surface Area (SASA)

To further characterize the conformational dynamics of the tau fibrils over the neuronal membrane and in the water medium, we have studied the radius of gyration (R*g*). The distribution of the radius of gyration for the tau structures in the neuronal membrane and the water medium are shown in Figure 5. The radius of gyration of the independent simulations are included in the Supplementary Information (Table S2 – S3). The straight filament structures show a larger deviation in the radius of gyration values in the water medium compared to the neuronal membrane. In comparison, the paired helical filaments show a more conserved conformational ensemble in the neuronal membrane and the water medium. This is in accordance with the results obtained in the earlier RMSF values which show that the PHF structure is more stable than the SF structure. The higher flexibility of the SF structure in turn leads to the increase in the radius of gyration (R*g*) compared to the PHF structure.

**Fig. 5.**
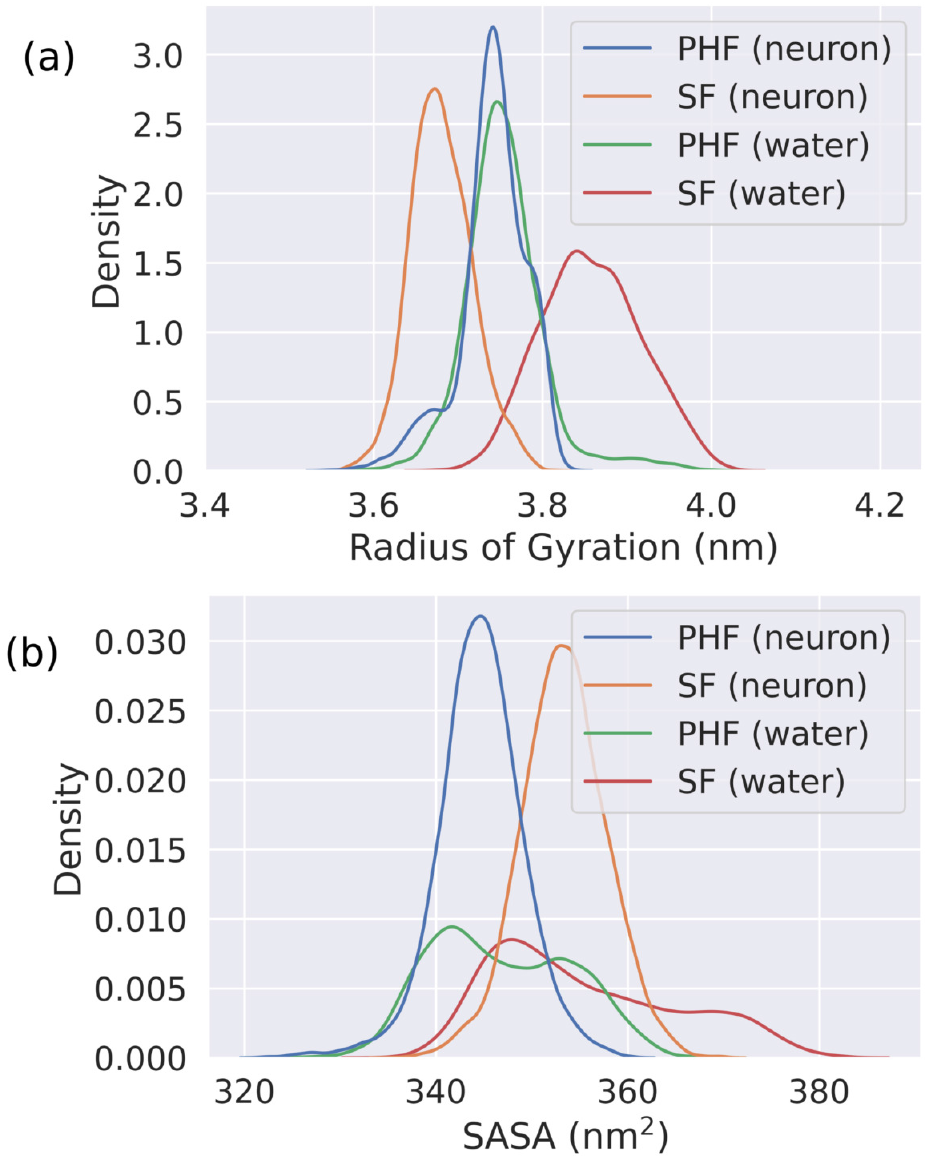
Distribution of (a) radius of gyration (R_*g*_) and (b) solvent accessible surface area (SASA in nm^2^) for the tau fibrils in the neuronal membrane and water medium.

The solvent accessible surface area (SASA) is calculated using a probe radius of 0.14 nm to trace the effect of neuronal membrane on the tau fibril polymorphs. The distribution of the SASA values are shown in Figure 5(b). The SASA values of the independent simulations are given in the Supplementary Information (Table S4 – S5). The SF structure is more soluble than the PHF structure in the presence of the neuronal membrane. The SF structure is also marginally more soluble in the water medium than the PHF structure. The distribution of the SASA values is broader in the water medium than in the neuronal membrane. The difference of the SASA values of SF structure in water and neuronal membrane are minimal with the mean value of SASA being 355.51 nm^2^ in water and 353.33 nm^2^ in neuronal membrane. The corresponding values for the PHF structures are 345.90 nm^2^ and 344.61 nm^2^ respectively, in the water medium and the neuronal membrane. Thus, the influence of the neuronal membrane on the SASA values compared to the pure water medium is negligible. The SF structure has higher SASA with respect to the PHF structure.

### 3.3 Dimer Interface and Cation-*π* contacts

The core residues of the PHF and SF structures are the same. The polymorphism arises from a difference in the dimer interface. The dimer interface of the PHF and SF fibrils is stabilized by a few specific amino acid residues. The important amino acid residues with the intra-residue distances between them are shown in Figure 6(a)-(b). The residues L315, K317, K321 are the primary residues at the SF dimer interface. In the PHF dimer interface, the saltbridge interactions between K331–E338 and the residues G334 play important role in stabilizing the fibril. The distance profiles between the dimer residues are shown in Figure 6. The cation – *π* interactions between the I308–Y310 residues play an important role in the tau stabilization and self-assembly as shown from the earlier experimental reports^54,55^. As a key residue of the steric zipper PHF6 fragment, Y310 plays an important role in tau fibrillization^7,56,57^. The cation – *π* contact is defined if the distance between the isoleucine and tyrosine residue is within 0.6 nm.

**Fig. 6.**
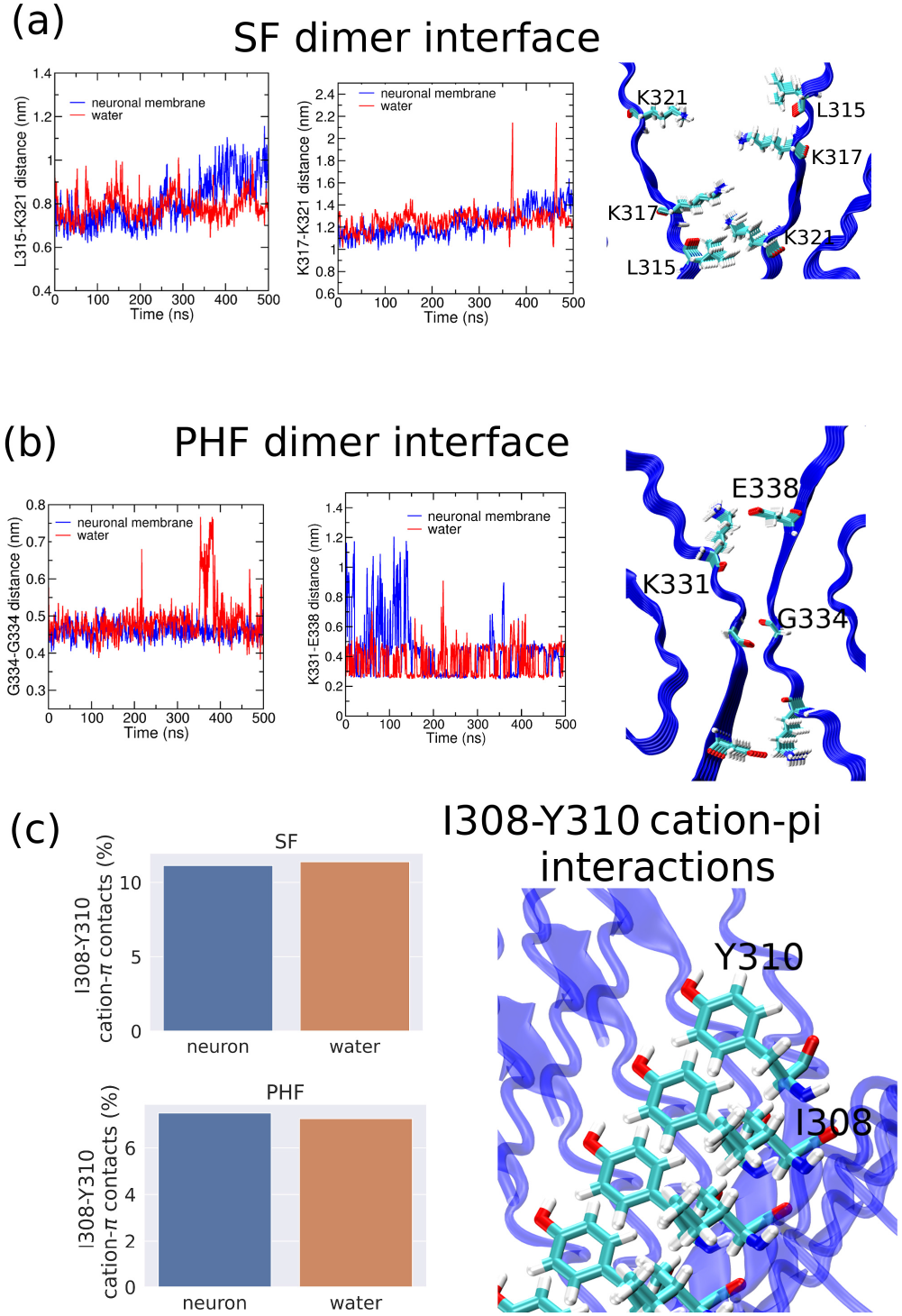
The primary amino acid residues at the dimer interface of the (a) SF and (b) PHF fibrils and (c) the cation – *π* interaction between I308–Y310.

Our results have shown that in the case of the SF dimer, the dimer interface consisting of L315–K321 has a distance of 0.935 nm in the cryo-EM structure. The average L315–K321 distance changes from 0.860 nm in the neuronal membrane to 0.783 nm in the water medium. The distance between K317– K321 also decreases in going from the neuronal membrane (1.31 nm) to the water medium (1.26 nm). The corresponding cryo EM distance in the pdb structure is 1.213 nm. Hence, the cryo EM distances for the interface residues change in the MD simulations both in the neuronal membrane and water medium. The L315–K321 distance decreases whereas, the K317–K321 distance increases from its value in the cryo EM structure.

In the PHF structures, G334–G334 distance in the cryo EM structure is 0.599 nm, whereas the distance in the neuronal membrane and water medium are 0.458 nm and 0.484 nm, respectively. That is, G334–G334 distance decreases both in the case of the neuronal membrane and the water medium. For the salt bridge interactions between K331–E338 residues, the distance between the 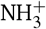 and COO− groups in the cryo EM structure is 0.693 nm which changes to 0.423 nm in the neuronal membrane and 0.371 nm in the water medium. Thus, the saltbridge interaction increases in both the neuronal membrane and the water medium. Hence, in the MD simulations in both the neuronal membrane and the water medium; the intra-residue distances in the dimer interface change in both the tau polymorphs. Moreover, the PHF dimer interface is strengthened by the decrease in the intra-residue distance in the neuronal membrane and the water medium compared to the cryo EM structure.

There is negligible change in the cation – *π* contacts between I308–Y310 residues in the neuronal membrane and water medium simulations among both the tau polymorphs. However, the cation – *π* contacts are lower in PHF structures and there is nearly a 40% decrease in the cation – *π* contacts in the PHF structures compared to the SF structure. Therefore, the cation – *π* interactions are relatively stronger in the SF structure resulting in their higher propensity for self assembly.

### 3.4 Secondary Structure Content

The molecular dynamics simulations of the IDPs shed light on the secondary structure content^30^. The choice of the appropriate force field along with the choice of the water model used in molecular dynamics simulations influence the secondary structure of the IDPs^58–60^. As shown from the earlier studies with regards to the A*β* monomer and fibril structures, secondary structure content is dictated by the nature of the interaction with the bilayer membranes^33,35,61,62^. In the neuronal membrane, the *β* - sheet structures are reduced in the case of SF structures whereas *β* -sheet content increases in the case of PHF structures as shown in Figure 7A. There is nearly a 8% decrease in the *β* -sheet content in case of PHF structures in the water medium compared to the neuronal membrane whereas in case of SF structure, the *β* -sheet content increases nearly 8% in going from neuronal membrane to the water medium. The cryo-EM structure of the tau fibril cover the residues V306–K311 in *β* 1, V313–C322 in *β* 2, N327–K331 in *β* 3, Q336–S341 in *β* 4, K343–K347 in *β* 5, R349–I354 in *β* 6, S356–V363 in *β* 7 and N368–F378 in *β* 8. Thus, these eight zones of *β* -sheet residues are studied using the DSSP utility in *gmx*. The residue-wise propensity of the secondary structures are shown in Figure 7B(a-d) for the tau structures in the neuronal membrane and aqueous medium. The decrease in the *β* -sheet content in the water medium for the PHF structure is seen prominently at the C-terminal regions. This decrease of the *β* -sheet content leads to the concomitant increase in the coil conformations. Similarly, in the SF structures, the *β* -sheet content is increased in the C-terminal region along with the increase of the coil conformations. It is to be noted that the C-terminal region is spanned by the *β* 8 region of the tau between N368–F378 residues in the cryo-EM structures which shows maximum changes in going from the neuronal membrane to the aqueous medium.

**Fig. 7.**
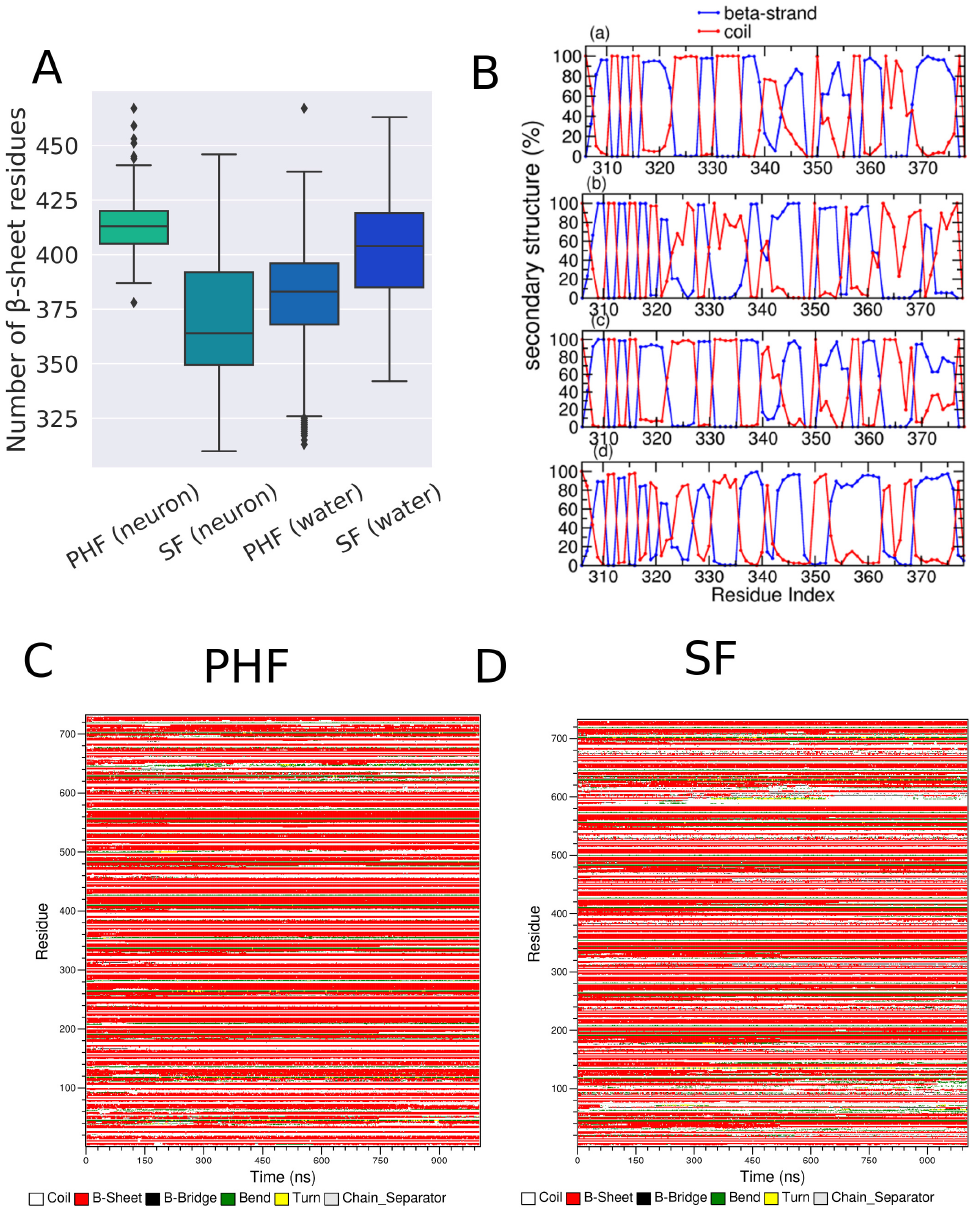
(A) The total number of *β* -sheet residues in the four systems, shown in a box and whisker plot. (B) (a)-(b) are the plots showing the average residue-wise secondary structure propensity in case of PHF and SF structures in neuronal membrane and (c)-(d) are the average secondary structure propensities for the PHF and SF structures in the water medium. (C)-(D) are the DSSP timelines for the PHF and SF structures in the neuronal membrane.

The experimental results show contrasting behavior regarding the secondary structure content on tau fibril binding to the lipid membranes. The tau K19 protein interacting with vesicles show the formation of helices in the R3–R4 domain^63^. The formation of the *α*-helix conformations might be observed at much larger timescales, or the absence of R1 region might be limiting the biophysical modeling. On the contrary, the hexapeptide fragment (PHF6) shows the formation of the *β* -sheet structures in complexation with the dissolved lipids containing negatively charged DMPG lipids^53^.

### 3.5 Number of Contacts

To characterize the fibril – lipid interactions, we have calculated the number of contacts per lipid molecule in the neuronal membrane which is shown in Figure 8(a)-(b). The contact is defined using a cutoff of 1.0 nm between the atoms of the fibril and the lipid molecules. The GM1 lipids show most number of contacts with PHF structure. The second highest contacts with PHF are observed for POPC lipids. In case of the PHF structure, the number of contacts between the fibril, and POPC and GM1 lipids are 64.93 and 72.61 respectively. PSM and POPS lipids have the least and second lowest number of contacts with the PHF fibril with the values of 10.19 and 13.17 respectively. The most number of contacts in SF structure are found in case of negatively charged GM1 lipids (53.61), followed by POPC lipids (57.80). The number of contacts for the zwitterionic PSM and negatively charged POPS are 18.94 and 17.08. Thus, the GM1 and POPC lipids preferentially interact with both the tau polymorphs in the neuronal membrane.

**Fig. 8.**
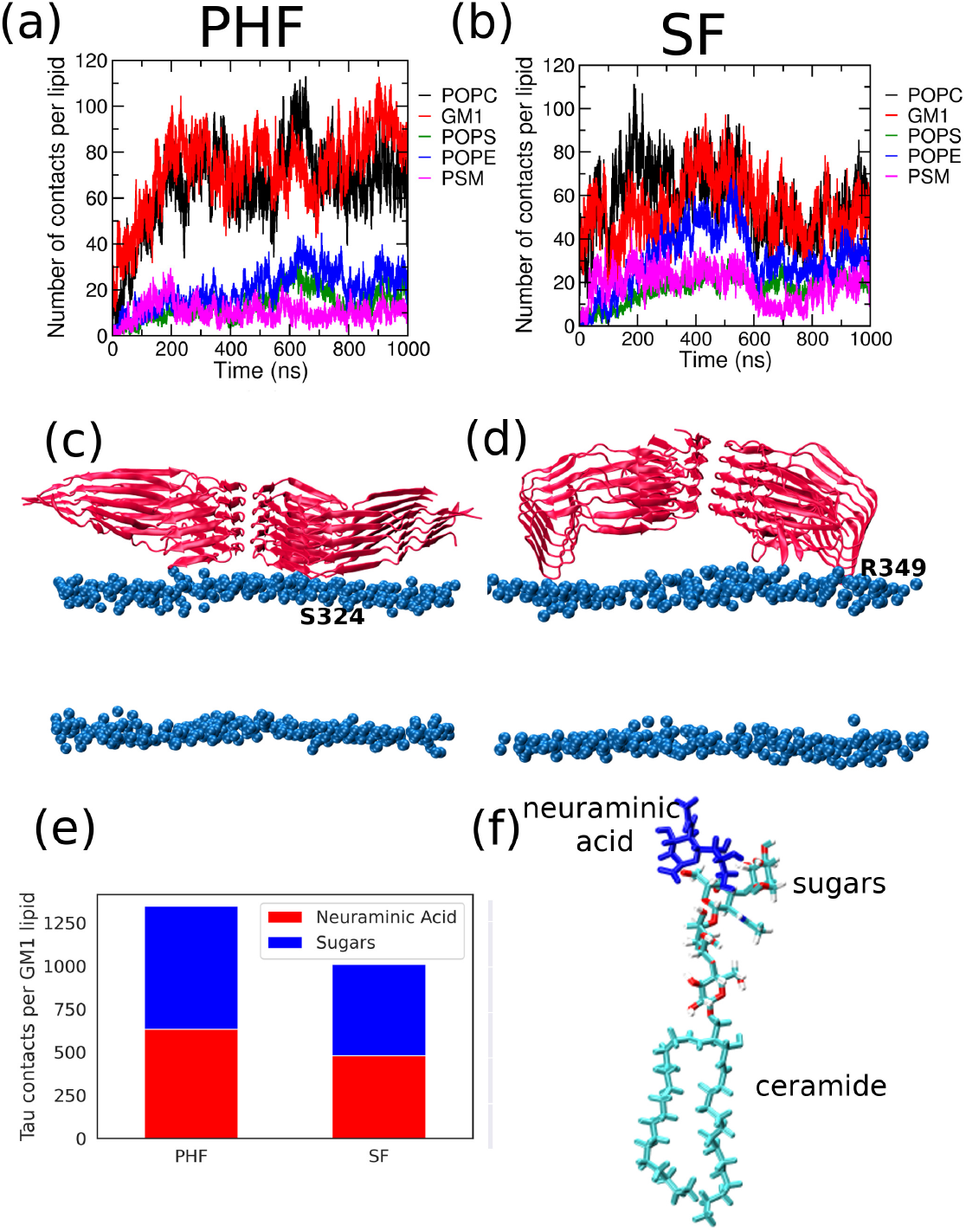
(a)-(b) are the number of contacts per lipid molecules for the PHF and SF structures. (c)-(d) are the most populated cluster for the tau fibril over the neuronal membrane obtained using the clustering algorithm. The protein is shown in new cartoon representation and the phosphorus headgroups are shown as blue VDW spheres. The interacting residues are marked in the figure for the two structures. (e) is the bar plot showing the number of contacts of the GM1 lipid groups with the fibrils per GM1 molecule. (f) show the individual groups within the GM1 lipids. The ceramide tail and neuraminic acid are shown in cyan and blue respectively.

To further resolve the interaction of the tau fibrils with the GM1 lipids, the number of contacts between the neuraminic acid and the sugar molecules were determined and are shown in Figure 8(e). The number of contacts for the neuraminic acid and the sugar groups are normalised with respect to the total number of GM1 lipids present in the two systems. The representative snapshot of GM1 lipid showing the sugar molecules and the neuraminic acid is given in Figure 8(f). The sugar molecules show only a small increase (nearly 10 %) in the number of contacts with the tau polymorphs compared to the neuraminic acid. The large number of contacts in case of the GM1 lipids are due to its protein localization ability through the formation of suitable carbohydrate-amino acid interactions. The GM1 lipids with their sugar groups can anchor the tau filaments over the membrane thus inducing higher number of contacts as shown in Figure 10(d)-(e). Our findings are also consistent with the earlier reports related to the role of GM1 lipids in the amyloid *β* and *α*-synuclein aggregation^64–67^.

The most populated structure generated from the simulation trajectories through clustering algorithm are shown in Figure 8(c)-(d). The PHF and SF structures are seen to interact with the neuronal membrane in a different manner. In the PHF structure, S324 is seen to interact with the phosphorus head group of the bilayer, whereas in the SF structure, positively charged R349 is found to interact with the phosphorus head group.

### 3.6 Hydrogen Bonding

To characterize the interactions between the tau fibril and the neuronal membrane, we have calculated the number of hydrogen bonds in all the systems. We have classified the hydrogen bonding according to the geometric criteria^68^. In a strong hydrogen bond, the hydrogen atom and the acceptor are separated by a distance less than 0.22 nm, and the angle made by the donor, hydrogen atom, and the acceptor is within the range 130° – 180°. The corresponding distance and angular range are 0.2 – 0.3 nm and 90° – 180°, respectively, for a weak hydrogen bond. We did not find strong hydrogen bonding in any of the systems.

The hydrogen bonding numbers in the studied systems are given in Table 2. The intrafibril hydrogen bonding stabilizes the fibril and helps conserve the *β* -sheet. Thus, the variation in the tau fibril hydrogen bonding among the two polymorphs and in the two media follows similar trend as the number of *β* -sheet residues. The number of intrafibril hydrogen bonds decreases in going from the neuronal membrane to the water medium in the case of PHF structure, whereas it increases in the case of SF structure. The changes in the number of intramolecular hydrogen bonds are small in both the systems (3.38% and 0.47% in the PHF and SF structures respectively). The fibril-GM1 hydrogen bonding is predominant which stabilizes the fibril-membrane interaction.

**Table 2.** Number of hydrogen bonds observed in the studied systems. Fibril – lipid hydrogen bonds have been normalised with respect to the number of individual lipid components.

|  | PHF |  | SF |  |
| --- | --- | --- | --- | --- |
|  | Neuronal membrane | Water | Neuronal membrane | Water |
| intra-fibril | 613.54 | 592.79 | 582.98 | 585.75 |
| Fibril – GM1 | 0.462 | – | 0.385 | – |
| Fibril – POPS | 0.215 | – | 0.360 | – |
| Fibril – POPC | 0.018 | – | 0.021 | – |
| Fibril – POPE | 0.028 | – | 0.062 | – |
| Fibril – PSM | 0.008 | – | 0.042 | – |

To further characterize the stability of the neuronal membrane, we calculated the intrafibril hydrogen bonding between the lipid molecules, which are shown in Table 3. The number of hydrogen bonds is averaged over all the lipid molecules. The pure membrane lipid molecules have less number of intra-lipid hydrogen bonds with respect to the tau fibril incorporated neuronal membranes. The intra-PSM hydrogen bonds are higher in number than the intra-POPS, intra-POPC and intra-POPE hydrogen bonds. This increase of hydrogen bonding within the PSM lipids in mixed membranes is also illustrated earlier^69^.

**Table 3.**
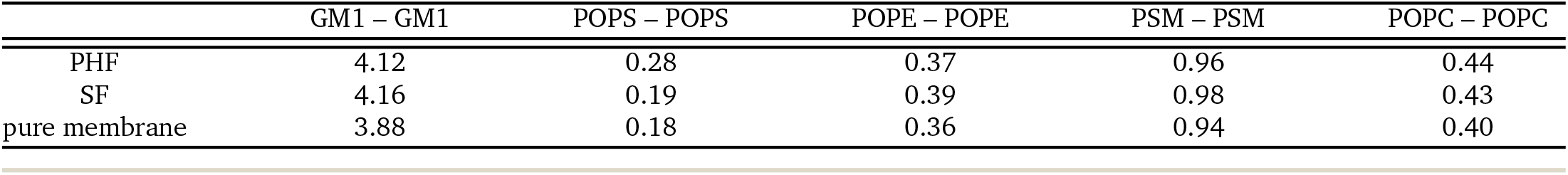
Number of intra-species hydrogen bonds among the lipid species for the different systems calculated per number of lipid molecules.

|  | GM1 – GM1 | POPS – POPS | POPE – POPE | PSM – PSM | POPC – POPC |
| --- | --- | --- | --- | --- | --- |
| PHF | 4.12 | 0.28 | 0.37 | 0.96 | 0.44 |
| SF | 4.16 | 0.19 | 0.39 | 0.98 | 0.43 |
| pure membrane | 3.88 | 0.18 | 0.36 | 0.94 | 0.40 |

### 3.7 Contact Map

The analysis of the contact map of the residues are shown in Figure 9 for the water medium and the neuronal membrane in case of both the polymorphs. The contact maps are calculated through the intra-residue distances with the distance being shown in a colorbar. The residue contact maps show the difference between the tau ensemble conformation in the two different environment. The structures are stabilized by the contact between far separated residues. The contact map show signature of the antiparallel *β* -sheet arrangement involving residues I308–F347 and residues K348–K370. These regions are dominated by the *β* -sheet regions in both the media. Additional V354–I372 contacts of the tau fibrils are less in the neuronal membrane compared to those in the water medium.

**Fig. 9.**
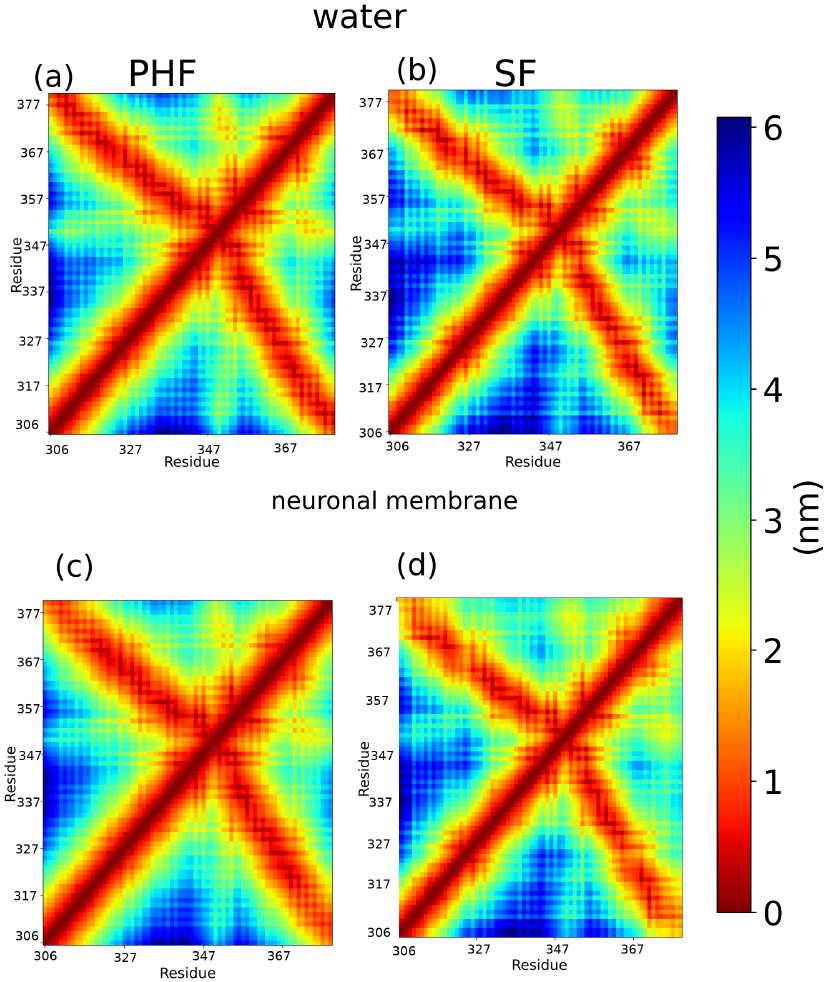
(a)-(b) are the residue-residue contact maps for the PHF and SF structures in the water medium. (c)-(d) are the contact maps in the neuronal membrane.

### 3.8 Bilayer Properties

It is difficult to study the effect of intrinsically disordered proteins on the membrane bilayer using conventional experimental techniques. Thus molecular dynamics simulations helps to elucidate the membrane protein interactions at an atomic resolution. To characterize the perturbations in the neuronal membrane due to the presence of the tau fibril over it, we have calculated the bilayer properties. The bilayer thickness in case of the tau incorporated membranes are similar to those of the pure membrane as shown in Table 4. The bilayer thickness is calculated by taking the headgroup to headgroup distance of the P-atoms in POPC, POPS, POPC, and PSM. The two dimensional bilayer thickness projected over the bilayer plane is shown in Figure 10(a)-(b). The area per lipid of the tau incorporated membranes is lower than that in pure membrane system. The change in bilayer thickness is negligible upon the incorporation of the tau fibrils. The cholesterol tilt angle is calculated taking the angle subtended by the C3–C17 carbon atoms of cholesterol with the bilayer normal.

**Table 4.**
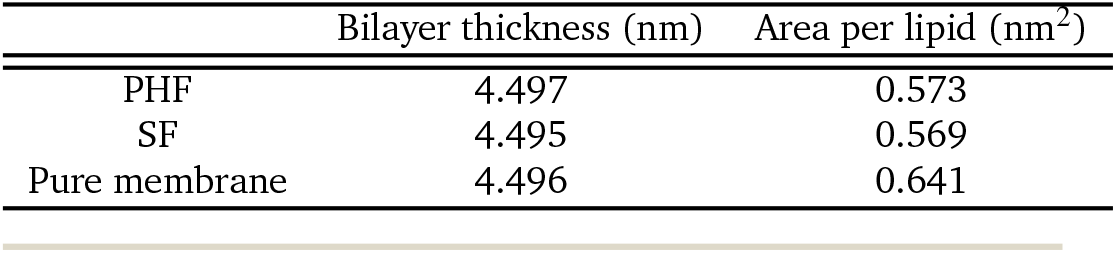
Bilayer thickness (nm) and the area per lipid (nm^2^) in the neuronal systems and the pure neuronal membrane.

**Fig. 10.**
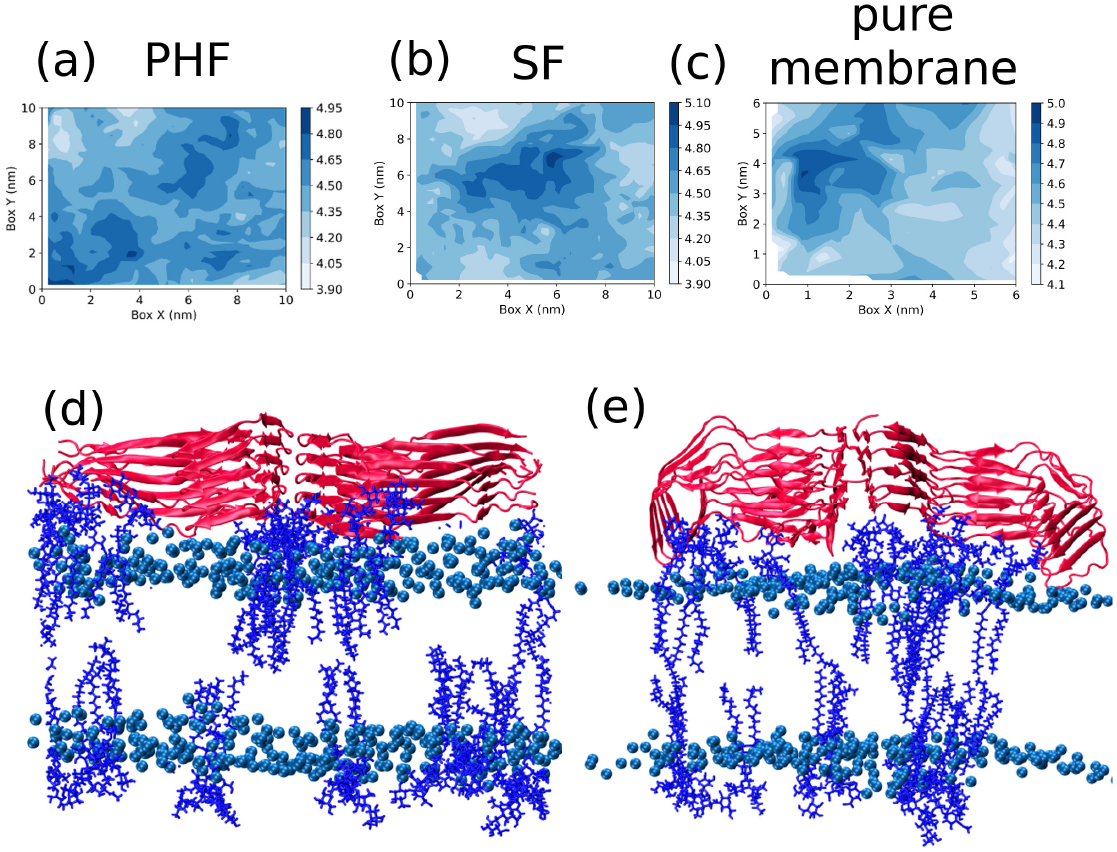
(a), (b), (c) are the two dimensional bilayer thickness projected over the bilayer plane for PHF, SF and pure membrane respectively. (d) and (e) are the final snapshots of the PHF and SF structures over the neuronal membrane respectively. The GM1 lipids are shown in blue licorice representation, phosphorus atoms are shown as VDW spheres and the tau fibril is shown in new cartoon representation.

The bilayer properties for the independent simulations are included in the Supplementary Information (Section 2.10). The area per lipid and the bilayer thickness are included in Table S8 and the two dimensional bilayer thickness are given in Figure S11 – S12 in the Supplementary Information. The cholesterol tilt angle distribution of the individual systems are included in the Supplementary Information (Figure S13). The bilayer thickness has more uniform distribution in the presence of tau fibrils. In comparison, the bilayer thickness for the amyloid *β* dimer over the neuronal membrane is calculated to be 4.65 *±* 0.03 nm by Fatafta *et al*.^35^. In general, we find that the bilayer properties like the bilayer thickness, cholesterol tilt angle show negligible change with the presence of the tau filaments. The relatively higher change in the bilayer properties is found in the area per lipid (APL). This result is found to be correlating with the earlier study concerning the amyloid *β* dodecamer and fibril over the neuronal membrane^70^.

### 3.9 Density Profile

The number density of the lipid components in the presence and absence of the tau polymorphs is shown in Figure 11. The density of the individual lipid molecules is projected over the bilayer plane and shown in a colorbar. Even without the tau fibrils the lipid molecules are distributed heterogeneously. GM1 tails (ceramide), POPS and PSM show distinct islands of high density. POPE and POPC lipids have a more uniform distribution. In case of POPE and POPC lipids, we observe large areas of high lipid distribution which results in uniform distribution of lipid molecules in the bilayer membrane.

**Fig. 11.**
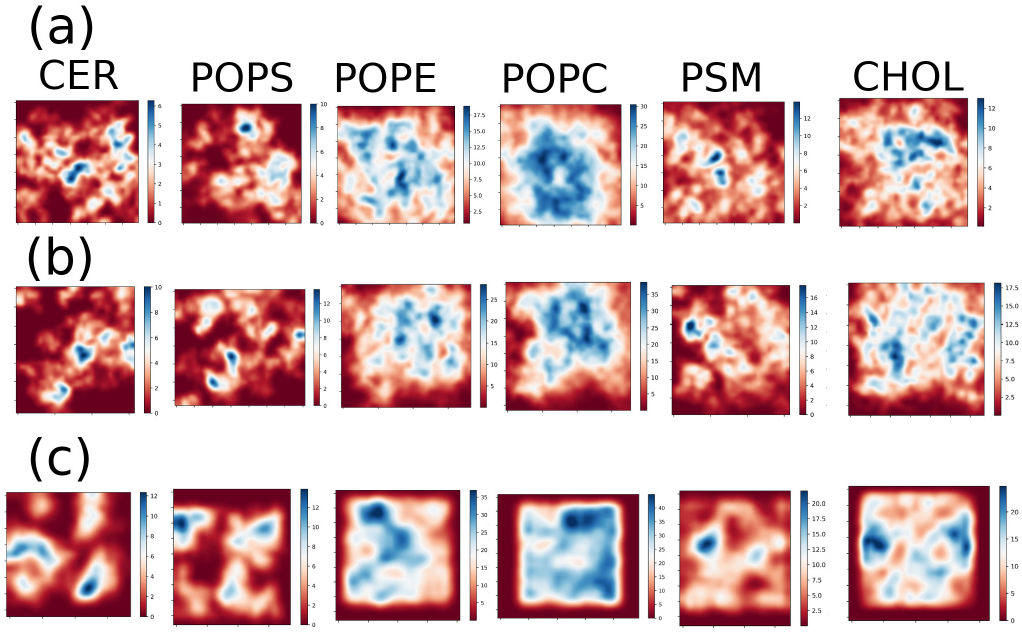
The density distribution of the lipid molecules projected over the bilayer plane in case of neuronal membrane with (a) PHF polymorph and (b) SF polymorph, and (c) pure neuronal membrane. The relative density is shown as a colorbar.

With the incorporation of tau fibril, the distribution of the lipid molecules is altered especially of the GM1 tails and POPS lipids. GM1 tails tend to cluster more in the absence of tau fibril. The distribution of cholesterol (CHOL) becomes even with the incorporation of tau fibril. The distribution of the lipid molecules is different in case of the PHF and SF structures. For the GM1 tails, the distribution is more widespread in the case of the PHF fibrils than the SF fibrils. The interaction of the tau fibrils with the neuronal membrane decreases the relatively broader distribution of the lipids. Hence, we conclude that the distribution of the lipid molecules over the bilayer changes in the presence of the tau polymorphs. Our results are consistent with the earlier studies related to the A*β* proteins over the neuronal membrane.^35,70^

## 4 Conclusions

Tau proteins have emerged as the important contributors to the tau pathogenesis. The impact of the tau fibrils on the neuronal membrane is pivotal to the understanding of the neuronal toxicity. We aim to understand the effect of tau fibril polymorphs on the neuronal membrane through all-atom molecular dynamics simulations. Control simulations have been performed on the pure neuronal membrane and the tau fibrils in water box. This is the first report of computational modeling aspiring to study the effect of tau fibrils on three or more lipid molecules forming the bilayer. The paired helical filament (PHF) in general show higher stability in the neuronal membrane compared to the straight filament (SF) structure as shown from the structural analyses. The tau binding over the neuronal membrane is predominantly due to the GM1 lipid molecules interacting with the tau fibril. The PHF structure is also found to be more compact in both the neuronal membrane and the water medium. The use of the entire cryo-EM structure for the initial configuration helps us to understand the changes that occur in the fibril interface of the PHF and the SF structures in aqueous solution and neuronal membrane. PHF and SF structures in the neuronal membrane show distinct differences in the number of *β* -sheet residues when compared with the simulations in the water medium. In the timescale of our simulations, we find that among PHF and SF, the PHF approaches closer to the neuronal membrane. There is a marked difference in the distribution of the lipid molecules in the presence of PHF and SF structures over the neuronal membrane. Overall, our study helps to describe the atomistic interactions responsible for the tau fibril binding to the neuronal membrane. Even though more extensive computational sampling starting with different initial conformations are required to unravel the fibril insertion into the membrane, the current study captures the disparate behavior of the tau polymorphs over the neuronal membrane. This in turn leads to pertinent questions like how the various set of polymorphs cause different tauopathies, which will require further studies despite a plethora of published literature on this field.

## Author Contributions

Unmesh D. Chowdhury : Conceptualization, Methodology, Software, Formal analysis, Investigation, Data curation, Visualization, Writing – Original draft

Arnav Paul : Software, Formal analysis, Data curation, Writing – Original Draft

B.L. Bhargava : Conceptualization, Methodology, Writing – Review & Editing, Resource, Supervision

## Acknowledgements

The authors gratefully acknowledge NISER Bhubaneswar for the computational resources.

