## Supplementary Information for "Interaction of the Tau fibrils with the neuronal membrane^†^"

### Contents

**1 Simulation Details** **S3**

1.1 Tau fibril - bilayer simulations . . . . . . . . . . . . . . . . . . . . . . . . . . S3

1.2 Fibril water simulations . . . . . . . . . . . . . . . . . . . . . . . . . . . . S4

1.3 Pure bilayer simulations . . . . . . . . . . . . . . . . . . . . . . . . . . . . S5

**2 Results and Discussions** **S7**

2.1 RMSD of the tau fibrils (1$\mu$s simulations) . . . . . . . . . . . . . . . . . . . . S7

2.2 RMSD of the tau fibrils for the independent simulations . . . . . . . . . . . . S7

2.3 RMSF of the tau fibrils for the independent simulations . . . . . . . . . . . . S8

2.4 Radius of Gyration ($R_g$) of the tau fibrils . . . . . . . . . . . . . . . . . . . . S9

2.5 SASA ($nm^2$) of the tau fibrils . . . . . . . . . . . . . . . . . . . . . . . . . S10

2.6 Secondary structure content of the tau fibrils . . . . . . . . . . . . . . . . . . S11

2.7 Distance of approach of the tau fibrils over the bilayer . . . . . . . . . . . . . S12

2.8 Positively charged residues on the tau fibril over the bilayer . . . . . . . . . . S13

2.9 The final structures for the PHF and SF fibrils . . . . . . . . . . . . . . . . . S14

2.10 Bilayer properties . . . . . . . . . . . . . . . . . . . . . . . . . . . . . . . S15

### 1 Simulation Details

#### 1.1 Tau fibril - bilayer simulations

The PHF and SF tau structures have been taken from the PDB ids 503L and 503T respectively. The fibrils were placed on top of the symmetric lipid bilayers comprising of POPC, POPS, PSM, CHOL and GM1 adding up to 550 and 652 lipid molecules in the systems with SF and PHF structures respectively. The CHARMM-GUI web interface was used to set the starting configuration of the fibril-neuronal membrane. Energy minimization was done to avoid the unfavourable contacts between the neighbouring atoms. Then, the six steps of equilibration were done according to the following steps:

Table S1: Energy minimization and the equilibration protocol, with the force constants (in kJ mol$^{-1}$ nm$^{-2}$)

| Step | Duration | Ensemble | BB restraint | SC restraint | LH restraint | LT restraint |
|---|---|---|---|---|---|---|
| Minimization | 5000 steps | — | 4000 | 2000 | 1000 | 1000 |
| Equilibration | 125 ps | NVT | 4000 | 2000 | 1000 | 1000 |
| Equilibration | 125 ps | NVT | 2000 | 1000 | 400 | 400 |
| Equilibration | 125 ps | NVT | 1000 | 500 | 400 | 200 |
| Equilibration | 500 ps | NVT | 500 | 200 | 200 | 200 |
| Equilibration | 500 ps | NVT | 200 | 50 | 40 | 100 |
| Equilibration | 500 ps | NVT | 50 | 0 | 0 | 0 |
| Production | 1 $\mu$s, 100 ns | NPT | 0 | 0 | 0 | 0 |

where BB - protein backbone, SC - protein sidechain, LH - lipid headgroups, LT - lipid tailgroups.

The final box lengths post equilibration were found to be 13.80 $\times$ 13.80 $\times$ 20.48 nm$^3$ in the case of PHF structure and 12.68 $\times$ 12.68 $\times$ 19.24 nm$^3$ in case of SF structure. The production runs were carried out for 1 $\mu$s and 100 ns$\times$2 overall for the fibril neuronal membrane simulations. The initial configurations of the tau fibril with the neuronal membrane along with the cuboidal box are shown in Figure S1.

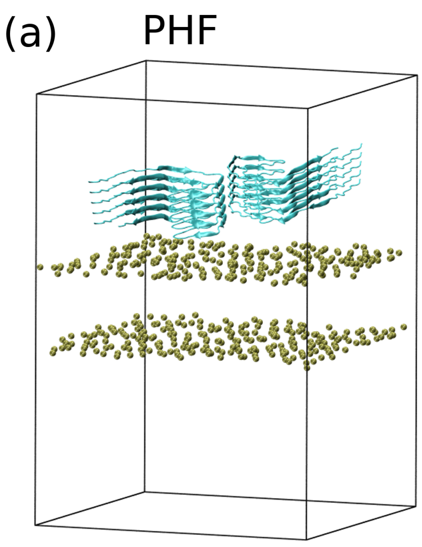


Figure S1: Starting configurations of (a) PHF and (b) SF structures with the neuronal membrane along with the cuboidal box. Only the phosphorus head groups of the bilayers are shown as VDW spheres for the ease of visualization.

#### 1.2 Fibril water simulations

The PHF and SF structures (PDB ids 503L and 503T) were immersed in a cubic box of water molecules. 0.15 (M) KCl salt was added to maintain the physiological concentration. The minimum distance between the edge of the box and the fibril was taken to be 1.5 nm and PBC has been applied in all the directions. The fibril and water have been modelled using CHARMM-36m and TIP3P water model respectively. Initally, the energy minimization was done restraining the protein backbone for 5000 steps. Then, the equilibration was done in the NVT ensemble at a temperature of 310K. The production runs were carried out in NPT ensemble for 500 ns or 100 ns as mentioned in the main manuscript. The timestep of 2 fs has been used and the bonds containing hydrogen atom were restrained using the LINCS algorithm. The long ranged cutoff of 1.2 nm was used with the Particle Mesh Ewald (PME) method. Isotropic pressure coupling was done through Parrinello-Rahman barostat ($\tau_p$=5 bar, compressibility=$4.5\times10^5$) and temperature coupling was done through Nosé-Hoover thermostat ($\tau$=1 ps)).

#### 1.3 Pure bilayer simulations

We have run the pure bilayer simulations without the tau fibril using less number of lipid molecules. 200 lipid molecules were taken as the control system for pure bilayer simulations. The bilayer comprised of 76 POPC molecules, 48 POPE molecules, 18 PSM molecules, 40 cholesterol molecules and 10 POPS molecules. Post equilibration the energy minimization was done through six steps as mentioned before in Table S1. The lipid molecules were modeled using the CHARMM-36 force field parameters along with TIP3P water model. The van der Waals and electrostatic interactions were cut off at 1.2 nm. The Parrinello-Rahman algorithm and semi-isotropic coupling method were applied to maintain the pressure at 1 bar with a relaxation time of 5 ps. The temperature was maintained at desired value (310 K) by the Nosé-Hoover algorithm and a relaxation time of 1 ps. The production runs were carried out for a total duration of 500 ns replicated twice to account for better sampling.

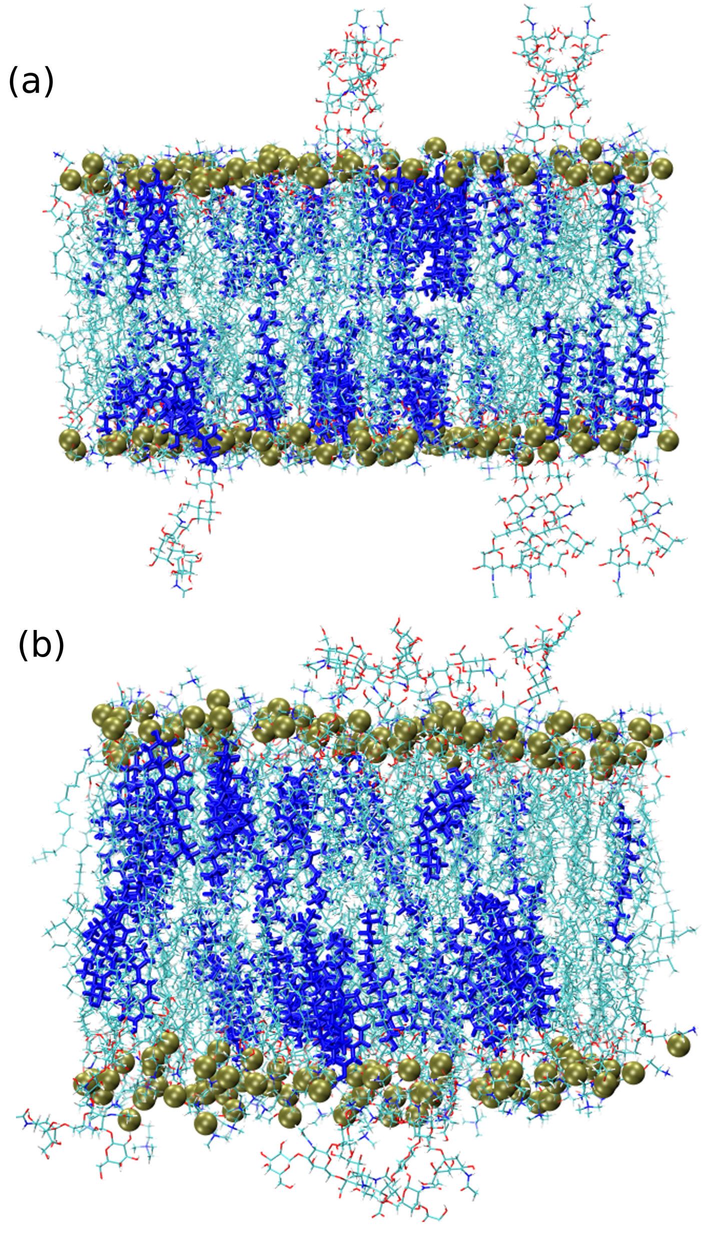


Figure S2: (a) The initial configuration of the neuronal membrane and (b) the final configuration of the neuronal membrane. The cholesterol molecules are shown in blue licorice representation, the phosphorus atoms as VDW spheres and the other lipid molecules are shown in line representation.

### 2 Results and Discussions

#### 2.1 RMSD of the tau fibrils (1$\mu$s simulations)

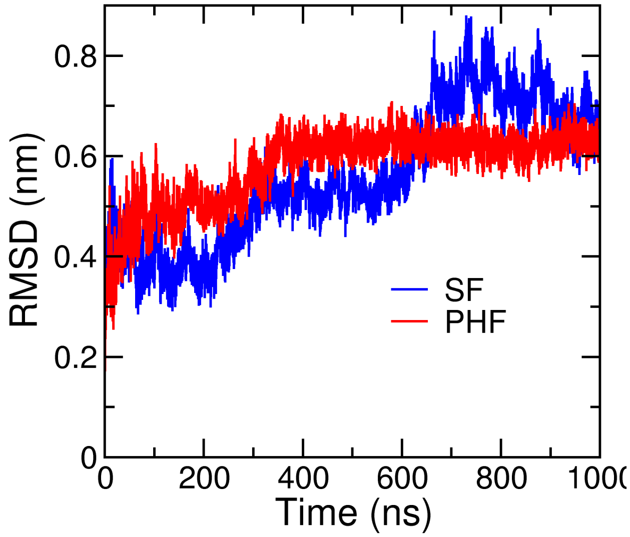


Figure S3: RMSD of the tau fibrils in neuronal membrane in 1$\mu$s simulation.

#### 2.2 RMSD of the tau fibrils for the independent simulations

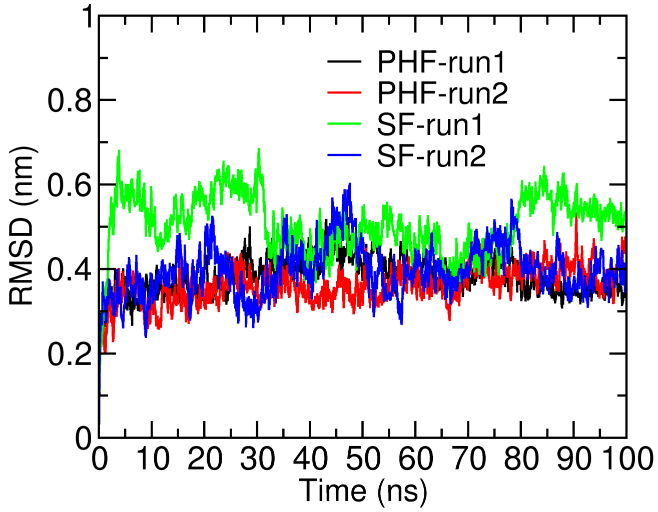


Figure S4: RMSD of the tau fibrils in neuronal membrane in independent simulations.

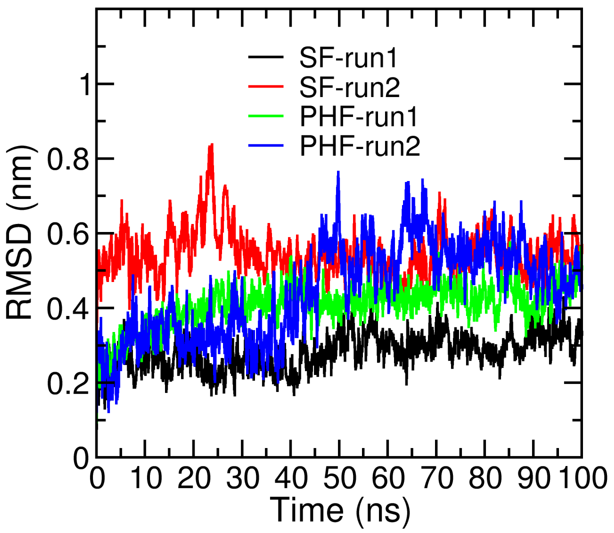


Figure S5: RMSD of the tau fibrils in water in independent simulations.

#### 2.3 RMSF of the tau fibrils for the independent simulations

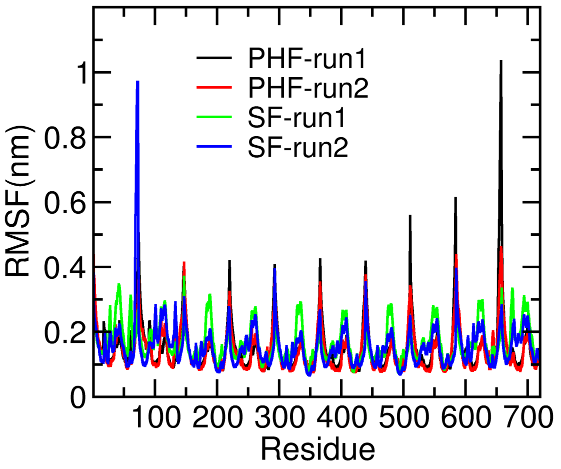


Figure S6: RMSF of the residues of the tau fibrils in the neuronal membrane obtained from the independent simulations.

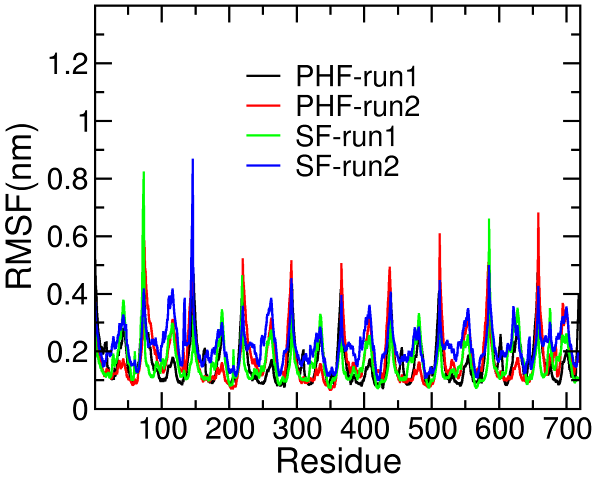


Figure S7: RMSF of the residues of the tau fibrils in the water box obtained from the independent simulations.

#### 2.4 Radius of Gyration ($R_g$) of the tau fibrils

Table S2: $R_g$ of the tau fibrils in the neuronal membrane obtained from the independent simulations.

| Systems | $R_g$ (nm) |
|---|---|
| SF-run1 | 3.64 ± 0.02 |
| SF-run2 | 3.65 ± 0.02 |
| PHF-run1 | 3.67 ± 0.02 |
| PHF-run2 | 3.63 ± 0.03 |

Table S3: $R_g$ of the tau fibrils in the water medium obtained from the independent simulations.

| Systems | $R_g$ (nm) |
|---|---|
| SF-run1 | 3.71± 0.02 |
| SF-run2 | 3.80 ± 0.03 |
| PHF-run1 | 3.67 ± 0.03 |
| PHF-run2 | 3.75 ± 0.03 |

#### 2.5 SASA ($nm^2$) of the tau fibrils

Table S4: SASA values of the tau fibrils in the neuronal membrane obtained from the independent simulations.

| Systems | SASA ($nm^2$) |
|---|---|
| SF-run1 | 341.91 ± 2.98 |
| SF-run2 | 343.23 ± 3.00 |
| PHF-run1 | 338.91 ± 3.60 |
| PHF-run2 | 335.67 ± 4.88 |

Table S5: SASA values of the tau fibrils in the water medium obtained from the independent simulations.

| Systems | SASA ($nm^2$) |
|---|---|
| SF-run1 | 341.48 ± 3.96 |
| SF-run2 | 346.47 ± 3.40 |
| PHF-run1 | 332.37 ± 3.39 |
| PHF-run2 | 335.34 ± 2.97 |

#### 2.6 Secondary structure content of the tau fibrils

Table S6: Average number of $\beta$-sheet residues and coil structures of the tau fibrils in the neuronal membrane obtained from the independent simulations.

| Systems | $\beta$-sheet | Coil |
|---|---|---|
| SF-run1 | 415.04 ± 10.68 | 251.55 ± 8.69 |
| SF-run2 | 434.14 ± 9.74 | 232.40 ± 7.84 |
| PHF-run1 | 417.511 ± 11.67 | 249.04 ± 9.97 |
| PHF-run2 | 418.43 ± 13.67 | 246.87 ± 11.21 |

Table S7: Average number of $\beta$-sheet residues and coil structures of the tau fibrils in the water medium obtained from the independent simulations.

| Systems | $\beta$-sheet | Coil |
|---|---|---|
| SF-run1 | 419.64 ± 8.04 | 248.01± 6.54 |
| SF-run2 | 427.43 ± 10.62 | 237.34 ± 10.06 |
| PHF-run1 | 407.88 ± 10.85 | 255.25 ± 11.15 |
| PHF-run2 | 385.26 ± 6.75 | 270.94 ± 6.47 |

#### 2.7 Distance of approach of the tau fibrils over the bilayer

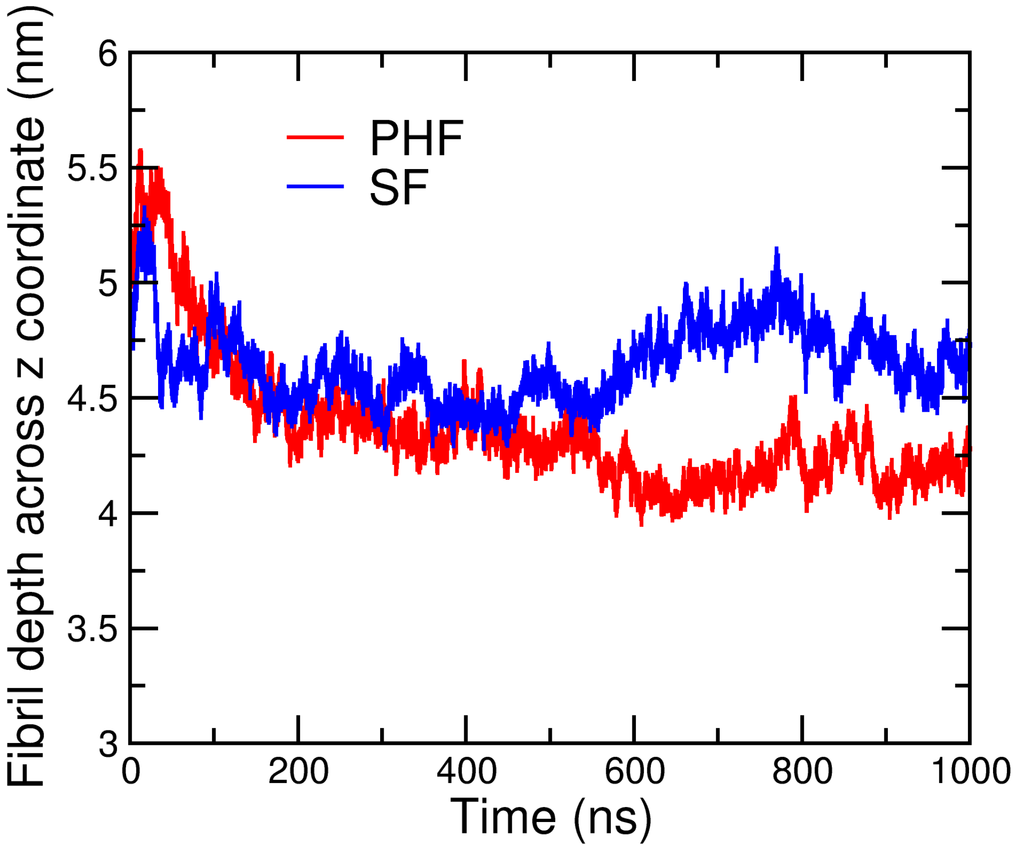


Figure S8: Distance of approach of the tau fibril over the neuronal bilayer along the Z-axis.

#### 2.8 Positively charged residues on the tau fibril over the bilayer

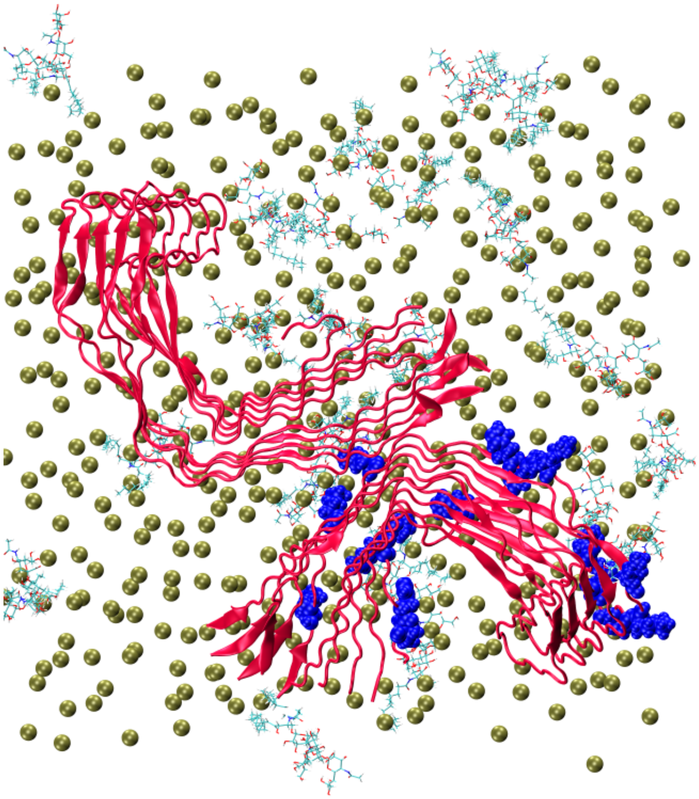

Figure S9: The positively charged residues are shown as blue VDW spheres. The sugar head groups of GM1 is shown in line respresentation, the phosphorus atoms of the bilayer is shown in VDW representation.

#### 2.9 The final structures for the PHF and SF fibrils

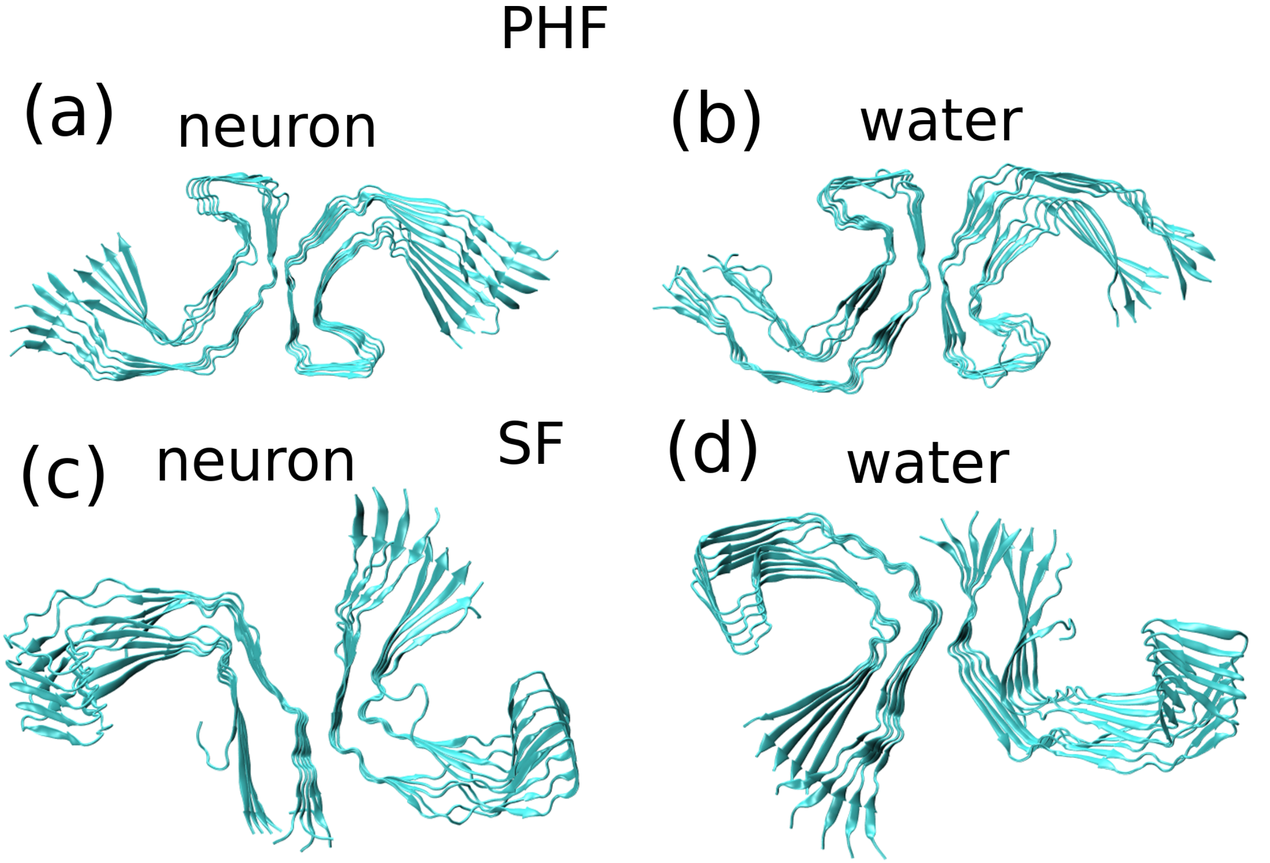


Figure S10: Final configurations of PHF structures in (a) neuronal membrane and (b) water medium and SF structures in (c) neuronal membrane and (d) water medium.

#### 2.10 Bilayer properties

Table S8: Bilayer thickness (nm) and area per lipids ($nm^2$) of the tau fibrils in the neuronal membrane obtained from the independent simulations.

| System | Thickness (nm) | Area per lipid ($nm^2$) |
|---|---|---|
| SF-run1 | 4.48 ± 0.06 | 0.58 ± 0.01 |
| SF-run2 | 4.46 ± 0.05 | 0.57 ± 0.01 |
| PHF-run1 | 4.46 ± 0.04 | 0.59 ± 0.01 |
| PHF-run2 | 4.47 ± 0.03 | 0.58 ± 0.01 |
| pure membrane | 4.49 ± 0.06 | 0.64 ± 0.01 |

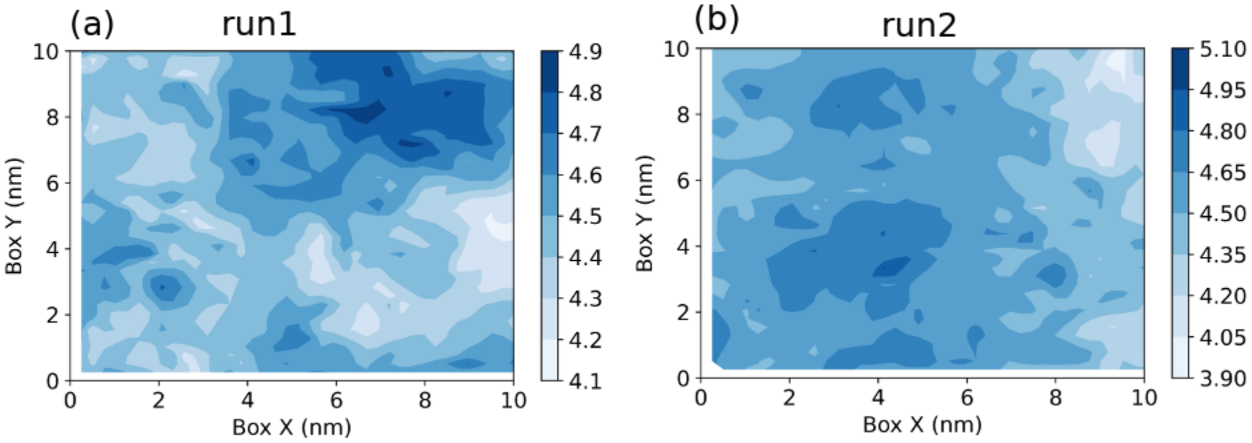


Figure S11: Two dimensional bilayer thickness in PHF – neuronal membrane system obtained from the independent simulations.

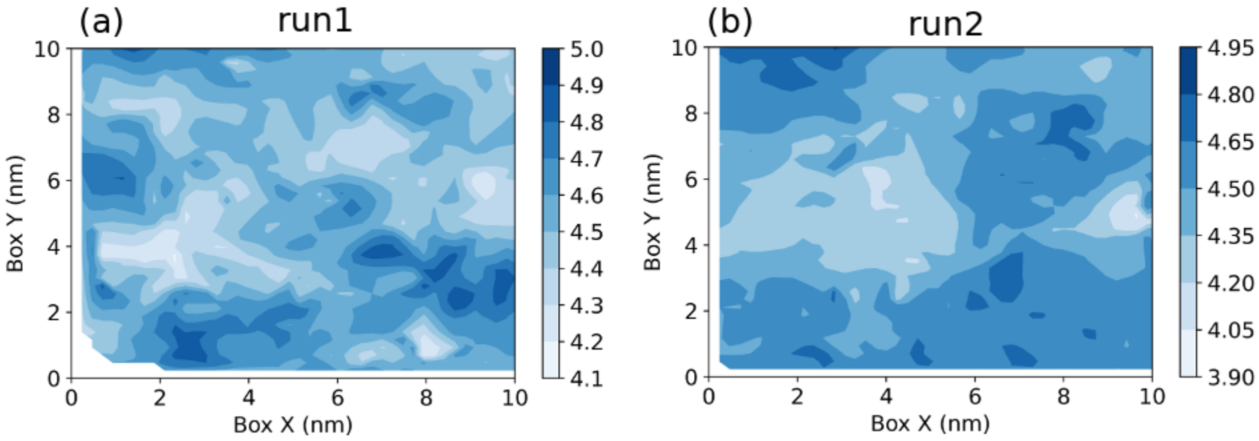


Figure S12: Two dimensional bilayer thickness in SF – neuronal membrane system obtained from the independent simulations.

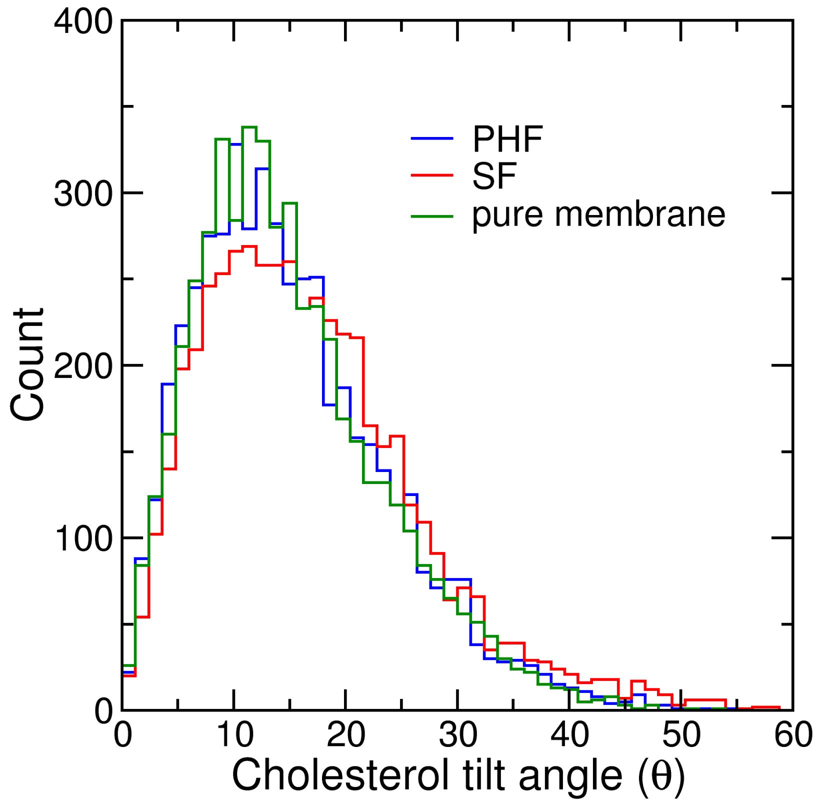


Figure S13: The tilt angle of the cholesterol with respect to the z-axis. The C3–C17 atoms are taken as the reference atoms. The tilt angle value for the pure neuronal membrane is $14.98 \pm 8.30$, the corresponding values for the PHF and SF structures are $14.91 \pm 8.33$ and $15.22 \pm 8.82$ respectively.
